# Ebola virus inclusion body formation and RNA synthesis are controlled by a novel domain of NP interacting with VP35

**DOI:** 10.1101/2020.04.06.028423

**Authors:** Tsuyoshi Miyake, Charlotte M. Farley, Benjamin E. Neubauer, Thomas P. Beddow, Thomas Hoenen, Daniel A. Engel

## Abstract

Ebola virus (EBOV) inclusion bodies (IBs) are cytoplasmic sites of nucleocapsid formation and RNA replication, housing key steps in the virus life cycle that warrant further investigation. During infection IBs display dynamic properties regarding their size and location. Also, the contents of IBs must transition prior to further viral maturation, assembly and release, implying additional steps in IB function. Interestingly, expression of the viral nucleoprotein (NP) alone is sufficient for generation of IBs, indicating that it plays an important role in IB formation during infection. In addition to NP, other components of the nucleocapsid localize to IBs, including VP35, VP24, VP30 and the RNA polymerase L. Previously we defined and solved the crystal structure of the C-terminal domain of NP (NP-Ct), but its role in virus replication remained unclear. Here we show that NP-Ct is absolutely required for IB formation when NP is expressed alone. Interestingly, we find that NP-Ct is also required for production of infectious virus-like particles and retention of viral RNA within these particles. Furthermore, co-expression of the nucleocapsid component VP35 overcomes deletion of NP-Ct in triggering IB formation, demonstrating a functional interaction between the two proteins. Of all the EBOV proteins only VP35 is able to overcome the defect in IB formation caused by deletion of NP-Ct. This effect is mediated by a novel protein-protein interaction between VP35 and NP that controls both regulation of IB formation and RNA replication itself, and which is mediated by a newly identified domain of NP, the “central domain” (CD).

**Importance:** Inclusion bodies (IBs) are cytoplasmic sites of RNA synthesis for a variety of negative sense RNA viruses including Ebola virus. In addition to housing important steps in the viral life cycle, IBs protect new viral RNA from innate immune attack and contain specific host proteins whose function is under study. A key viral factor in Ebola virus IB formation is the nucleoprotein, NP, which also is important in RNA encapsidation and synthesis. In this study, we have identified two domains of NP that control inclusion body formation. One of these, the central domain (CD), interacts with viral protein VP35 to control both inclusion body formation and RNA synthesis. The other is the NP C-terminal domain (NP-Ct), whose function has not previously been reported. These findings contribute to a model in which NP and its interactions with VP35 link the establishment of IBs to the synthesis of viral RNA.

## Introduction

Ebola virus (EBOV) causes severe, often fatal hemorrhagic disease in humans and is currently receiving enhanced attention due to a large recent outbreak in the Democratic Republic of Congo during late 2018 to early 2020. Promising vaccination development is underway, but efforts to better understand EBOV virology and pathogenesis so that additional approaches to prophylaxis and therapeutic discovery can be developed are still required (1-4).

Along with EBOV and the related Marburg virus (MARV), several well characterized negative strand RNA viruses, including vesicular stomatitis virus (VSV), respiratory syncytial virus (RSV), rabies virus (RABV), human parainfluenza virus (HPIV), Nipah virus (NiV), and human metapneumovirus (hMPV) carry out key replication steps in specialized cytoplasmic compartments referred to as inclusions, inclusion bodies (IBs) or Negri bodies (in the case of RABV) (5-19). In general, IBs are complex sites of viral RNA synthesis and contain viral proteins required for this process. In addition, IBs co-localize with specific host proteins whose roles in formation, maintenance and/or function of IBs are under study (19-25). Typically Ibs lack an organizing outer membrane and in some cases their formation is driven by liquid phase separation, a physical consequence of specific viral protein properties (26, 27). Evidence from RABV, RSV and EBOV suggests a dynamic relationship between IBs and virus-induced stress granules or specific stress granule proteins, and that this relationship regulates the innate immune response to infection (20, 23, 28). IBs are also thought to present a physical barrier to protect their RNA contents from innate immune attack. Precisely how IBs come into existence, the exact molecular processes they support during RNA synthesis, how they interfere with innate immunity and how they pass their contents along for further viral maturation are all incompletely answered questions that have garnered recent scrutiny.

IBs within EBOV infected cells are initially small and located near the endoplasmic reticulum, but become more widespread as infection progresses, and many increase in size (17, 29, 30). They contain viral proteins NP, VP35, VP40, VP30, VP24 and L, which are involved in various aspects of positive and negative sense RNA synthesis, nucleocapsid assembly and function, and viral maturation (30-32). Indeed, intact viral nucleocapsids have been visualized within EBOV IBs (18, 29, 33). The nucleoproteins (NP) of EBOV and MARV are key players in initiating IB formation, and cells ectopically expressing NP as the only viral factor contain “NP-induced IBs”, demonstrating that NP is sufficient for IB formation (10, 29, 34-37). In contrast, other viral nucleocapsid proteins fail to exhibit this behavior when individually expressed (34, 35, 38, 39), although there are somewhat conflicting data regarding the L polymerase in this regard. NP is a multifunctional 739 aa protein that is the most abundant component of the viral nucleocapsid, and in addition to triggering IB formation, its roles include RNA packaging, acting as a co-factor for RNA synthesis carried out by the viral polymerase L, and nucleocapsid assembly (32). A second viral protein, VP35, is also a required co-factor for EBOV and MARV RNA synthesis and is important in nucleocapsid function and assembly (40, 41). VP35 associates with NP and L, is found in IBs when co-expressed with NP, and one of its functions is to act as a bridge between NP and L in the formation of productive replication complexes (34, 35, 42). Physical interactions between VP35 and NP have been observed that involve both the N-terminal and C-terminal regions of NP (34, 41, 43-46), and these have been directly implicated in supporting viral RNA synthesis. Recently VP35 was shown to possess NTPase and helicase-like activities, which are proposed to support RNA remodeling during synthesis (47). VP35 also has well-documented anti-interferon (IFN) activity (48, 49).

Previously we reported the crystal structure of the C-terminal domain of NP (NP-Ct) from EBOV and the corresponding proteins from Taï Forest virus (TAFV) and Bundibugyo virus (BDBV) (50-52). NP-Ct is highly conserved across filoviruses and assumes a novel tertiary fold structure (50-53). Whereas activities carried out by the N-terminal domain of NP (aa 1-412; see Figure 1 and legend) have been well characterized and include RNA binding, NP-oligomerization, and physical association with VP35 and L, the activities of NP-Ct have remained a mystery. In this report, we demonstrate two novel and redundant functions of NP that control IB formation. One of these is carried out by NP-Ct, which we observe also plays a separate novel role in production of infectious transcription and replication-competent virus-like particles (trVLPs). The other IB-controlling function of NP is located within a previously uncharacterized region of the protein that spans amino acid positions 481-500, and is responsible for binding to the interferon inhibitory <u>d</u>omain (IID) of VP35. Importantly, we find this region of NP (the “central domain”; CD) to be crucially important not only for IB formation, but also for viral RNA synthesis. Together these findings reveal new activities for NP in several key viral replication steps and add to the complexity of viral RNA synthesis and IB dynamics that may potentially be exploited for small molecule inhibitor discovery.

**Figure 1.**
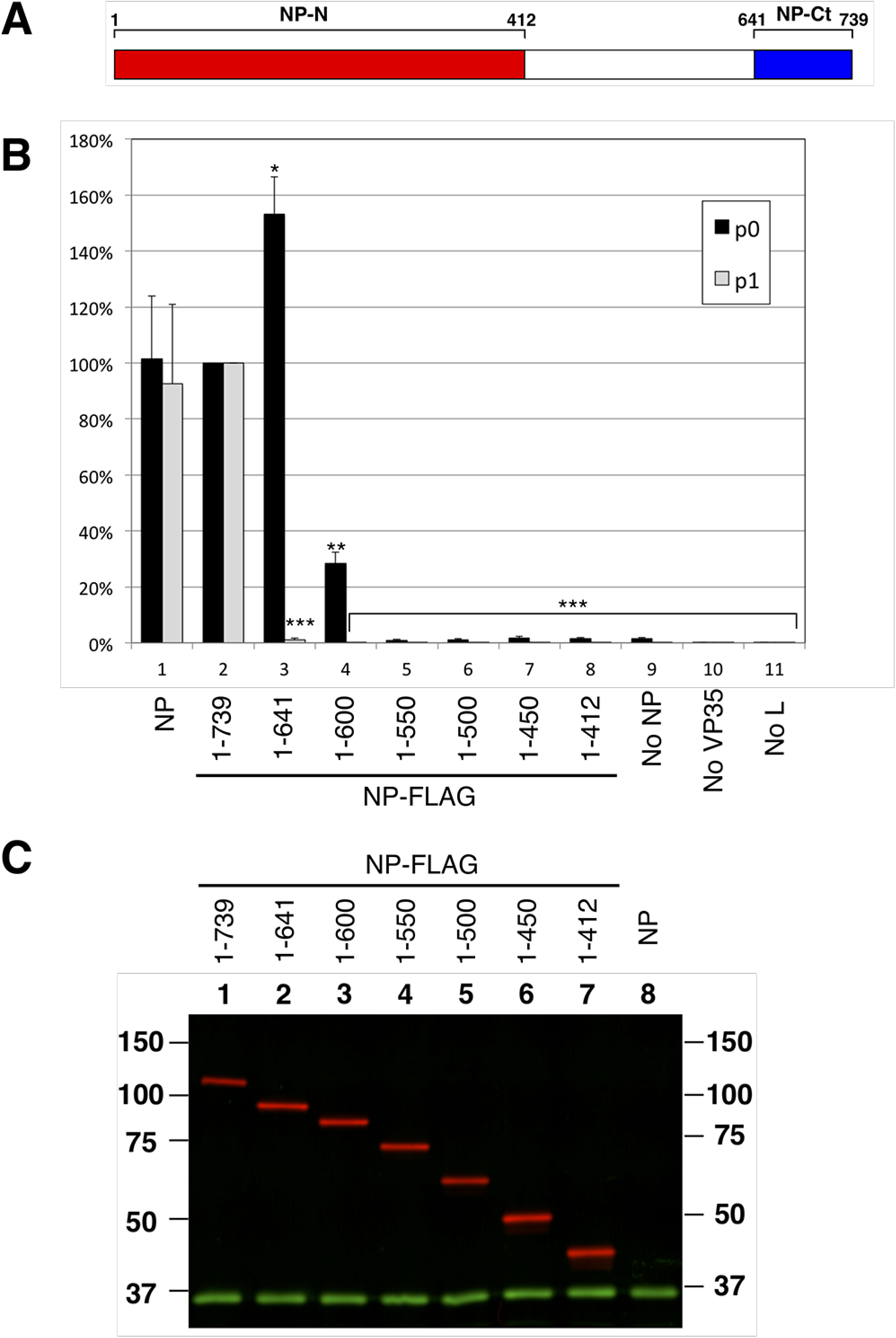
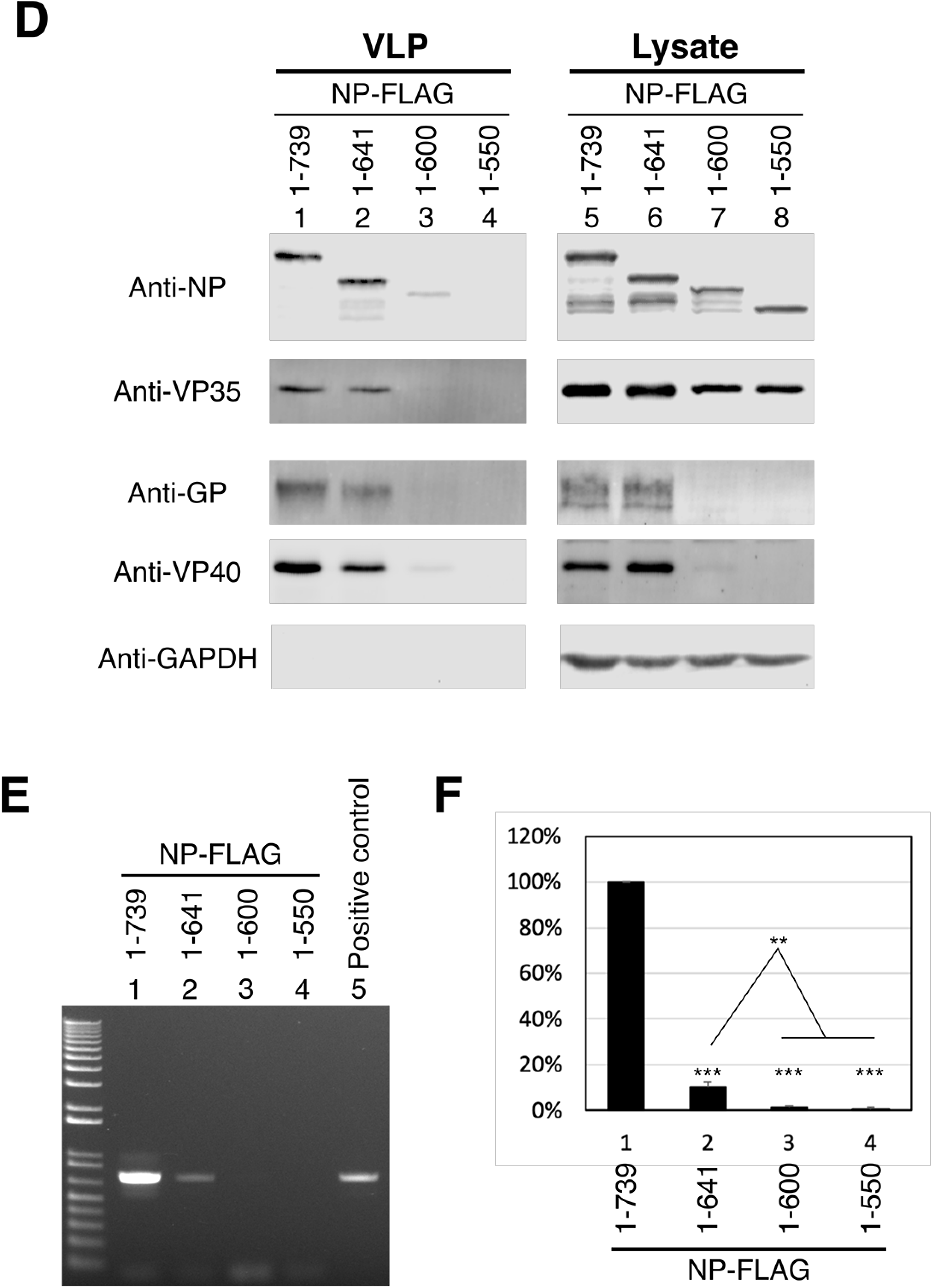
NP-Ct is required for infectious VLP production. A: NP primary structure. The precise definition of the N-terminal domain is subject to interpretation based on sequences included in constructs used for structural and biochemical studies (43, 44, 51, 66), but is labeled here as aa 1-412 based on sequence conservation. NP-Ct domain definition is based on Dziubanska et al. (51). B: trVLP assay (p0 cells) of NP deletion mutants. Indicated NP-FLAG constructs were transfected into 293T/17 cells. As described in the text, recipient p1 cells were supplied with wild-type NP. Data are averages of independent biological triplicates. Error bars represent the SD of the triplicates. Marks above the bar indicate: * *p*<0.05, ** *p*<0.01, *** *p*<0.001 to the corresponding NP-flag samples based on Student’s t-tests. Untagged NP (column 1) showed no statistically significant difference from the corresponding NP-flag samples. C: Western blot of lysates from p0 cell transfectants. Lysates were separated by SDS-PAGE, blotted and probed with anti-FLAG (red) or anti-GAPDH (green) antibody. D: Western blot analysis of purified trVLPs and corresponding lysate from p0 transfectants. Indicated NP-FLAG constructs were transfected into 293T cells (p0) along with all other trVLP components. trVLPs were purified from the supernatant as described in Materials and Methods. trVLPs and corresponding cell lysates were analyzed by western blot with the indicated antibodies. Anti-GAPDH was used as loading control for the lysate samples and as specificity control for the purified trVLPs. E: RT-PCR quantification of genomic (negative strand) RNA from the indicated, isolated trVLPs. Isolated RNA was subjected to RT-PCR and quantified using linear range amplification standards as described in Materials and Methods. Positive control indicates a PCR product of a minigenome DNA template. F: Quantitation of RT-PCR. PCR products were subjected to agarose gel electrophoresis and quantified with the Molecular Imager XRS system. Means and standard deviations of biologically triplicate experiments are shown. Asterisk *** indicates *p*<0.001 to the NP(1-739) sample, ** indicates *p*<0.01 between NP(1-641) and NP(1-600) or NP(1-550) based on Student’s t-test.

## Results

### NP-Ct is required for production of infectious VLPs but not for transcription or RNA replication

As illustrated in Figure 1A, NP-Ct spans amino acids 641-739, which corresponds to a region of high sequence conservation among *ebolavirus* species (51). To investigate its function, a series of deletion mutants were tested in the trVLP assay (54-56). In this assay, viral proteins VP40, VP24 and GP are expressed from a tetracistronic minigenome (MG), which also expresses Renilla luciferase as a reporter. All other viral proteins (NP, VP35, VP30 and L) are supplied individually by transfection. The transfected “p0” cells are competent for transcription, RNA replication and production of infectious VLPs containing newly replicated and encapsidated MGs. trVLPs can be recovered from the p0 supernatant and used to infect “p1” cells, which will produce new MGs and trVLPs if they also are supplied by transfection with plasmids encoding NP, VP35, VP30 and L. Importantly, we expressed NP deletion mutants in p0 cells but provided wild-type NP in p1 cells, allowing determination of whether p0 cells produced infectious trVLPs whose replication could be supported in p1 cells. Also, in p0 cells the pCAGGS-NP expression plasmid was replaced with a pCAGGS-NP-FLAG construct (and mutant derivatives), which we found to support trVLP activity equally to untagged NP (Figure 1B). Full length and all mutant NP proteins were expressed at very similar levels as shown in Figure 1C.

Deletion of the C-terminal 139 amino acid residues in NP(1-600) reduced reporter activity down to ∼28% of the full length protein in p0 cells, which is consistent with previous results (36) and is due to loss of a VP30 binding site (aa 600-615/617) that controls RNA synthesis, as described (57, 58). Mutant NP(1-550) or mutants with larger C-terminal deletions had less than 2% of wild-type reporter activity in p0 cells, with no statistically significant difference from a control lacking NP altogether, indicating a severe defect in transcription and/or RNA replication. In contrast, precise deletion of NP-Ct in NP(1-641) had no deleterious effect on reporter gene expression in p0 cells, indicating that NP(1-641) fully supports transcription and RNA replication. Strikingly however, when p0 cell supernatants from cells expressing NP(1-641) were used to infect p1 cells, reporter activity was only 1.0% of the wild-type, even though the p1 cells expressed wild-type NP supplied by transfection. These data demonstrate that despite wild-type levels of transcription and replication in p0 cells expressing NP(1-641), the p0 supernatants contained virtually no infectious trVLPs. To understand this infectivity defect, trVLPs were isolated from p0 supernatants and analyzed for protein and negative strand viral RNA content using our previously published methods (56). Interestingly, trVLPs isolated from p0 cells expressing either wild-type NP or NP(1-641) contained equal amounts of NP, indicating that the NP-Ct deletion did not result in a defect in NP incorporation into trVLPs. Furthermore, these trVLPs also contained wild-type levels of VP35, GP and VP40, demonstrating that they were indeed assembled trVLPs (Figure 1D lanes 1 and 2). However, expression of NP(1-600) or (NP1-550) did not produce intact trVLPs (Figure 1D lanes 3 and 4), consistent with the fact that the p0 cells expressing these proteins also did not express GP or VP40 (Figure 1D lanes 7 and 8), nor did they efficiently produce infectious VLPs as shown in Figure 1B. Importantly, trVLPs containing NP(1-641) contained significantly decreased amounts (10%) of genomic RNA compared with wild-type VLPs, which clearly correlates with their infectivity defect (Figures 1E and 1F). This indicates that NP-Ct has an important role in genomic RNA incorporation or stability within the trVLP’s nucleocapsid, and as such supports our conclusion of a novel function for this domain.

### NP-Ct is required for formation of inclusion bodies

As previously reported, NP from EBOV or MARV forms NP-induced IBs even in the absence of other viral proteins or viral RNA (10, 17, 29, 30, 34). To determine the role of NP-Ct in this process, HuH-7 cells were transfected with various NP-FLAG deletion constructs (Figure 2A) and stained with an anti-FLAG antibody (Figure 2B). Full length NP localized in IBs as expected, but NP(1-641), precisely lacking NP-Ct, clearly distributed throughout the cytoplasm and no IBs were observed. Also all other C-terminal deletion mutants of NP lacking NP-Ct, namely 1-600, 1-550, 1-500, 1-481 and 1-450, didn’t localize in IBs (Figure 2B). Expression of NP-Ct by itself as NP(640-739) was not sufficient for IB formation, and also NP(410-739) did not localize to IBs, indicating that additional N-terminal domains are also required. Deletion of the N-terminal aa 1-24 region, part of which is required for NP oligomerization and virus replication (43, 44) had no effect on IB formation, indicating that this region is dispensable for triggering IB formation. Together these results demonstrate that NP-Ct is an essential element for NP-induced IB formation.

**Figure 2.**
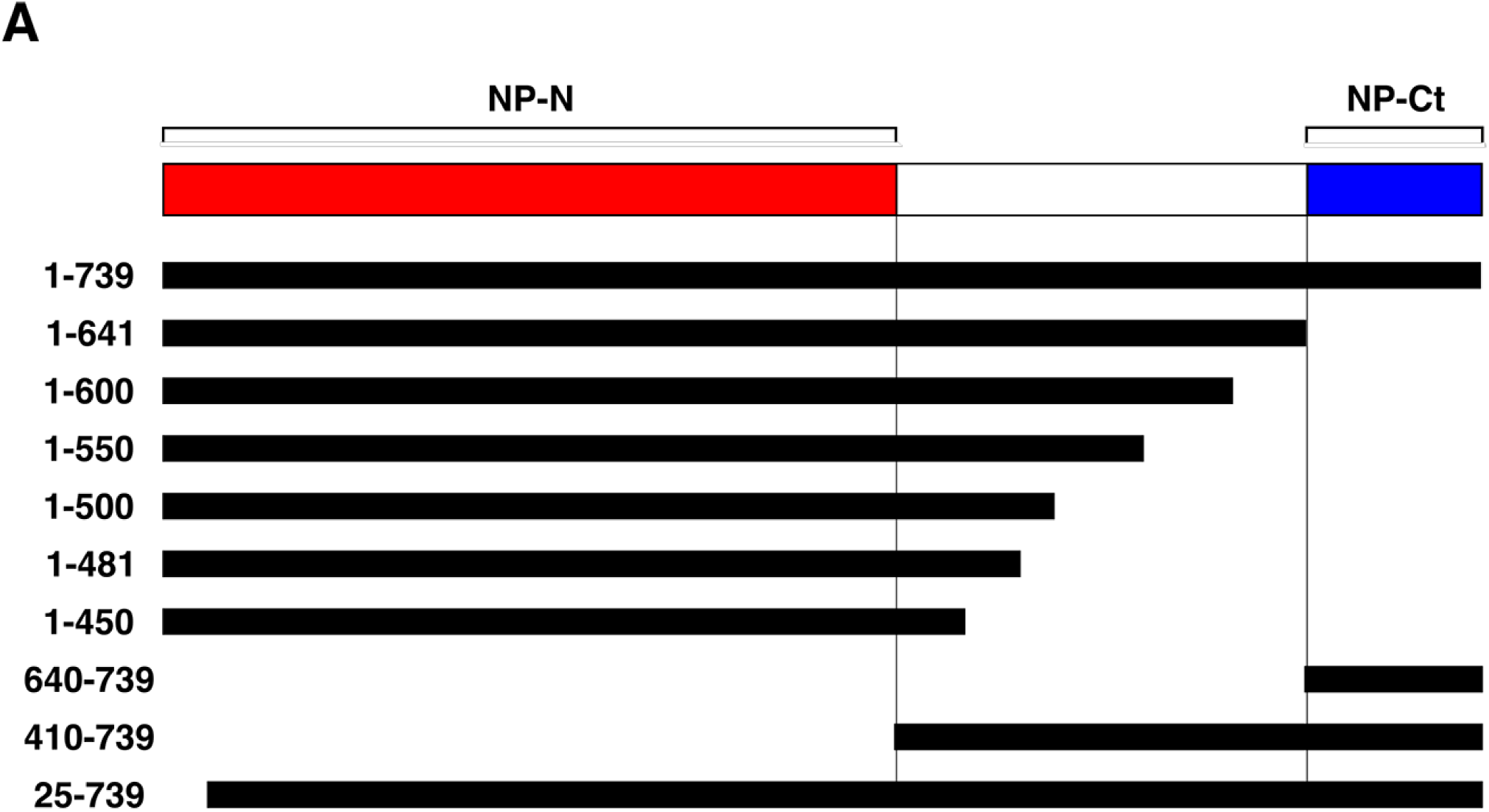
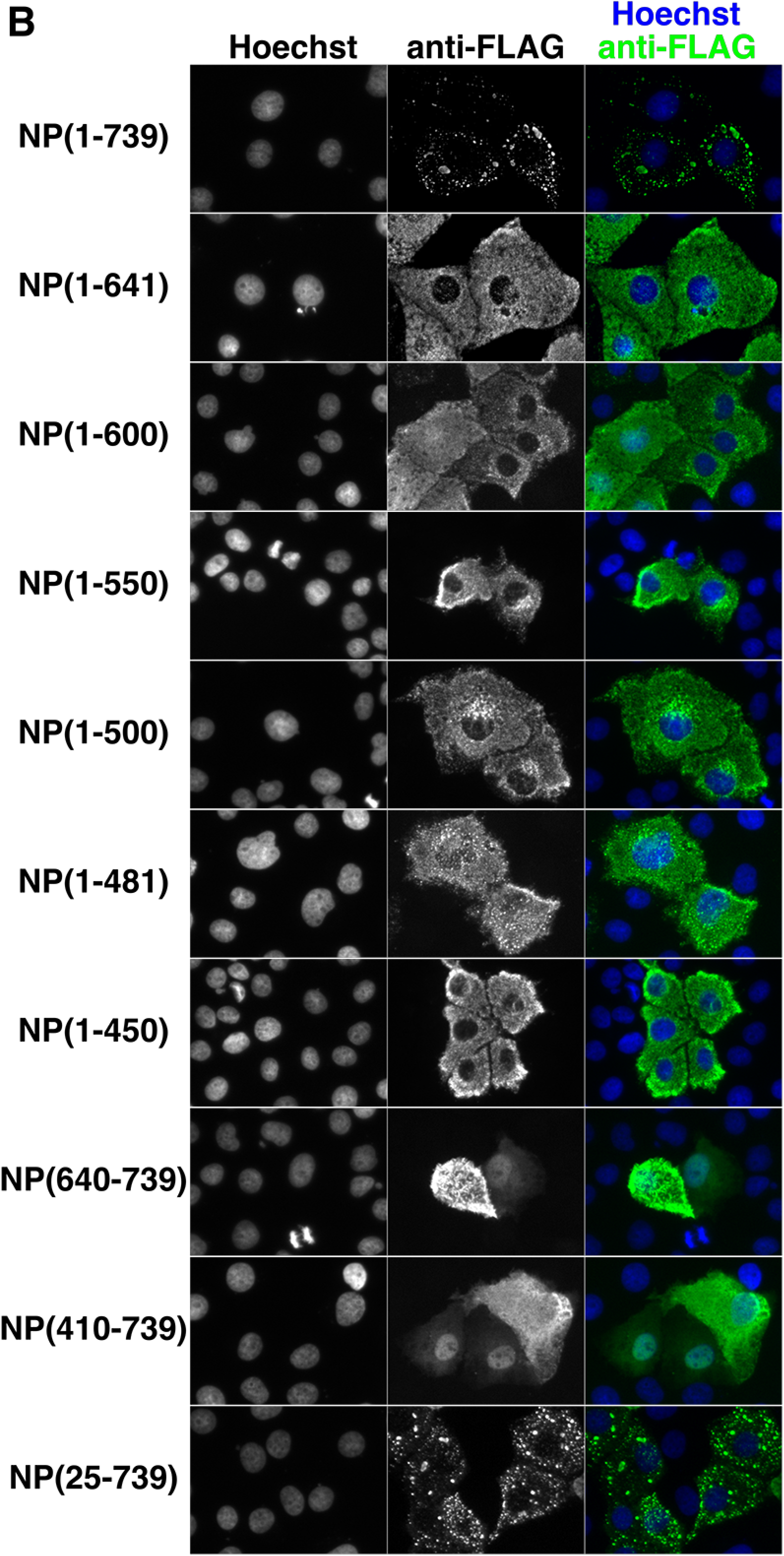
NP-Ct is required for IB formation. A: Structure of NP deletion mutants. Each construct contains a C-terminal FLAG-tag. B: HuH-7 cells were transfected with the indicated constructs and stained with anti-FLAG antibody and Hoechst 33342 dye after 48 hours. All the constructs lacking the NP-Ct failed to localize in IBs.

### VP35 specifically complements deletion of NP-Ct

We wondered if the role of NP-Ct in IB formation might be linked to its requirement for production of infectious VLPs as presented in Figure 1. Accordingly, we examined the localization of mutant NP(1-641) in the context of cells expressing all other trVLP assay components. Under these conditions NP(1-641) clearly localized in IBs (Figure 3A 2nd row), in complete contrast to its behavior when expressed alone (Figure 2). This indicated that one or more components of the trVLP system could complement the mislocalization of NP(1-641). Next, we systematically omitted each of the trVLP expression plasmids to identify which plasmid is necessary for complementing mislocalization of NP(1-641). As shown in Figure 3A, only omission of the VP35 expression plasmid, but not of any other trVLP system component, resulted in mislocalization of NP(1-641). To confirm that VP35 is indeed necessary and sufficient to complement the NP-Ct deletion, each plasmid of the trVLP system was transfected individually along with NP(1-641), as shown in Figure 3B. The VP35 expression plasmid was the only one that complemented mislocalization of NP(1-641).

**Figure 3.**
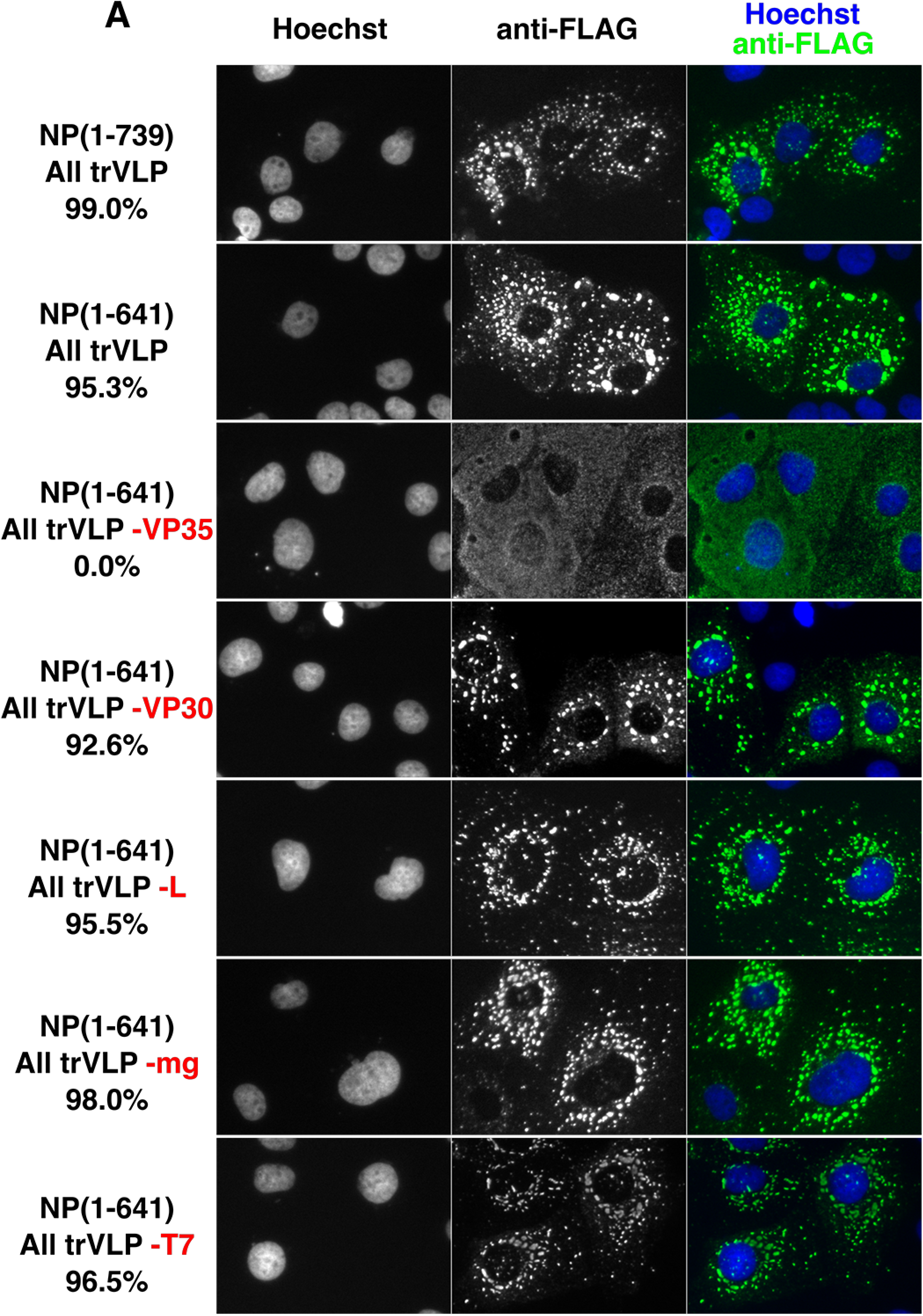
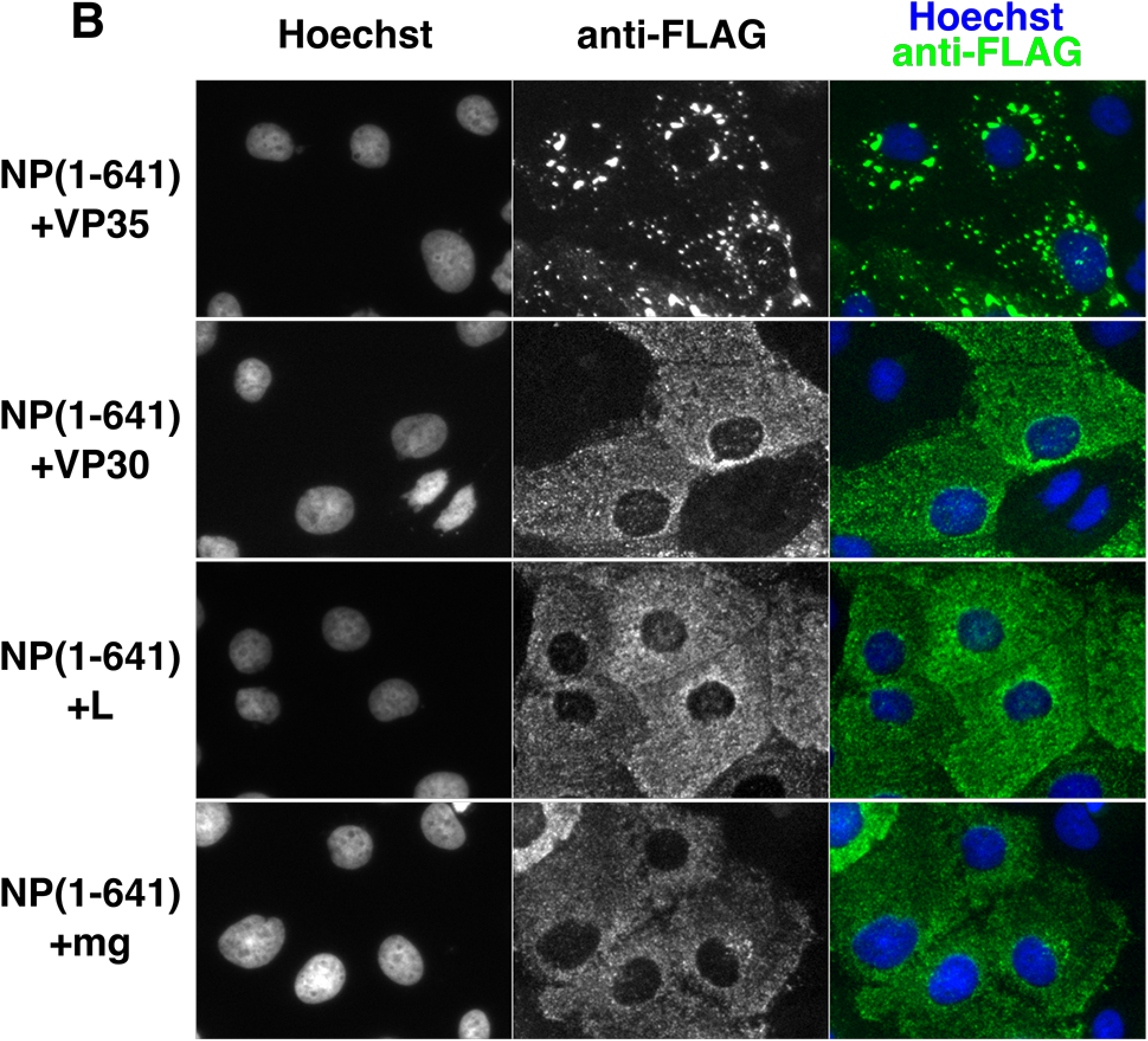
VP35 specifically complements deletion of NP-Ct. A: FLAG-tagged NP(1-739) or NP(1-641) was transfected along with other trVLP components, i.e. expression plasmids for VP35, VP30, L, T7 polymerase, a tetracistronic minigenome (mg) expressing VP24, GP and VP40, and firefly luciferase, as indicated, and immuno-stained with anti-FLAG antibody. Individually omitted constructs are indicated in red. B: NP(1-641) was co-transfected with each of the indicated individual plasmids of the trVLP system.

To avoid the potential problem of individual transfected cells receiving less than the full complement of plasmids, we used a lipid-based transfection with transfection complexes containing numerous plasmid copies. Given that the molecular weight of the transfected plasmids was between 3.5 and 7×10^6 g/mol, and that there were ∼1×10^6 cells per well at the time of transfection, this means that for each cell there were >10,000 plasmid copies available for transfection even of the lowest amounts (i.e. 75 ng). Thus, if a cell took up a transfection complex, it was very unlikely that plasmids of a single type would be absent from the complex. More importantly, our data clearly show that when we omitted the VP35 plasmid, we observed a dramatic change in phenotype, which we did not observe in any of the other cases (Figure 3). To confirm the data in Figure 3 indeed represent the typical phenotype of transfected cells, we counted 200+ cells per sample for each biological replicate, comparing NP in the inclusion bodies with overall NP-positive cells. In every case except the sample omitting the VP35 plasmid, the IB phenotype was >90%, of stained cells, and in the sample omitting VP35, it was 0%. These results demonstrate that VP35 is necessary and sufficient for NP(1-641) localization to IBs. VP35 is a co-factor for EBOV RNA synthesis and also has anti-interferon (IFN) activity (31). Since VP35 fails to trigger IB formation on its own (10, 34), these data also demonstrate that in the context of NP(1-641) expression, only two proteins, NP and VP35, are sufficient for IB formation. The data also show that mislocalization of NP(1-641) was likely not the cause of failure to produce infectious VLPs in p0 supernatants (Figure 1), because in cells that included NP(1-641) and all other trVLP components including VP35, intact IBs were observed, with proper NP(1-641) localization. This strongly suggests the presence of dual, separate functions in NP-Ct, one involved in infectious VLP production and the other controlling IB formation.

### The 481-500 region of NP is required for IB formation and for association with VP35

Because NP(1-641) localized in IBs when co-expressed with VP35, additional constructs were created to test which NP region is specifically required for IB formation in the presence of VP35, as presented in Figure 4A. FLAG-tagged C-terminal deletion mutants of NP were co-transfected with VP35. Full length NP(1-739) as well as all C-terminal deletion mutants up to NP(1-500) co-localized efficiently with VP35 in IBs, but neither NP(1-481) nor NP(1-450) did so. This indicates that the NP 481-500 region is required for IB localization when co-expressed with VP35, and explains the complementing activity of VP35 toward NP(1-641). However, as shown in Figure 1B, deletion mutants 1-550 or further C-terminal deletions showed trVLP activity of less than 1%, and the same was true for an additional mutant, NP(1-481)flag (data not shown). Therefore, the NP-VP35 interaction is not sufficient to support replication and transcription in the EBOV trVLP assay.

**Figure 4.**
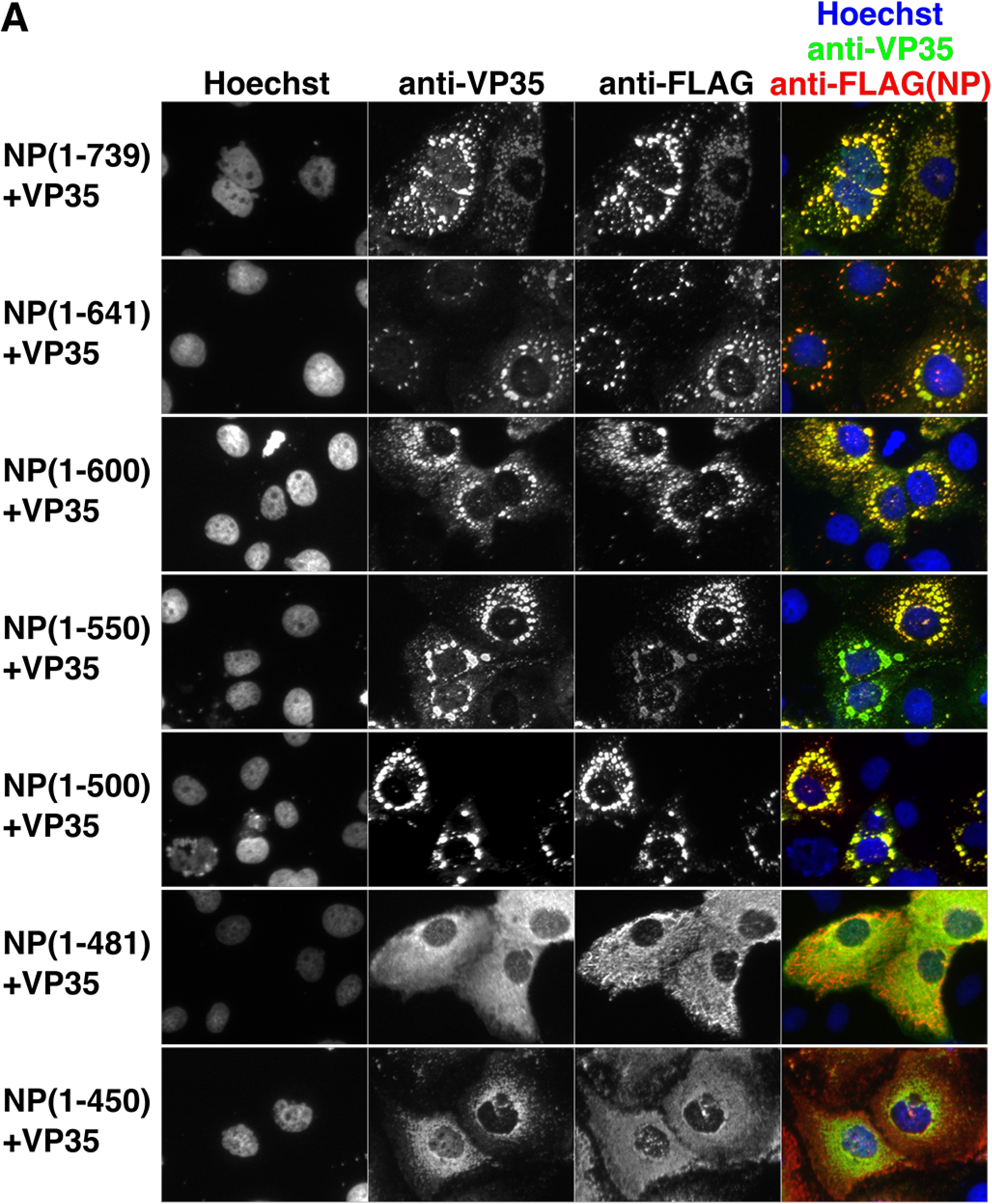
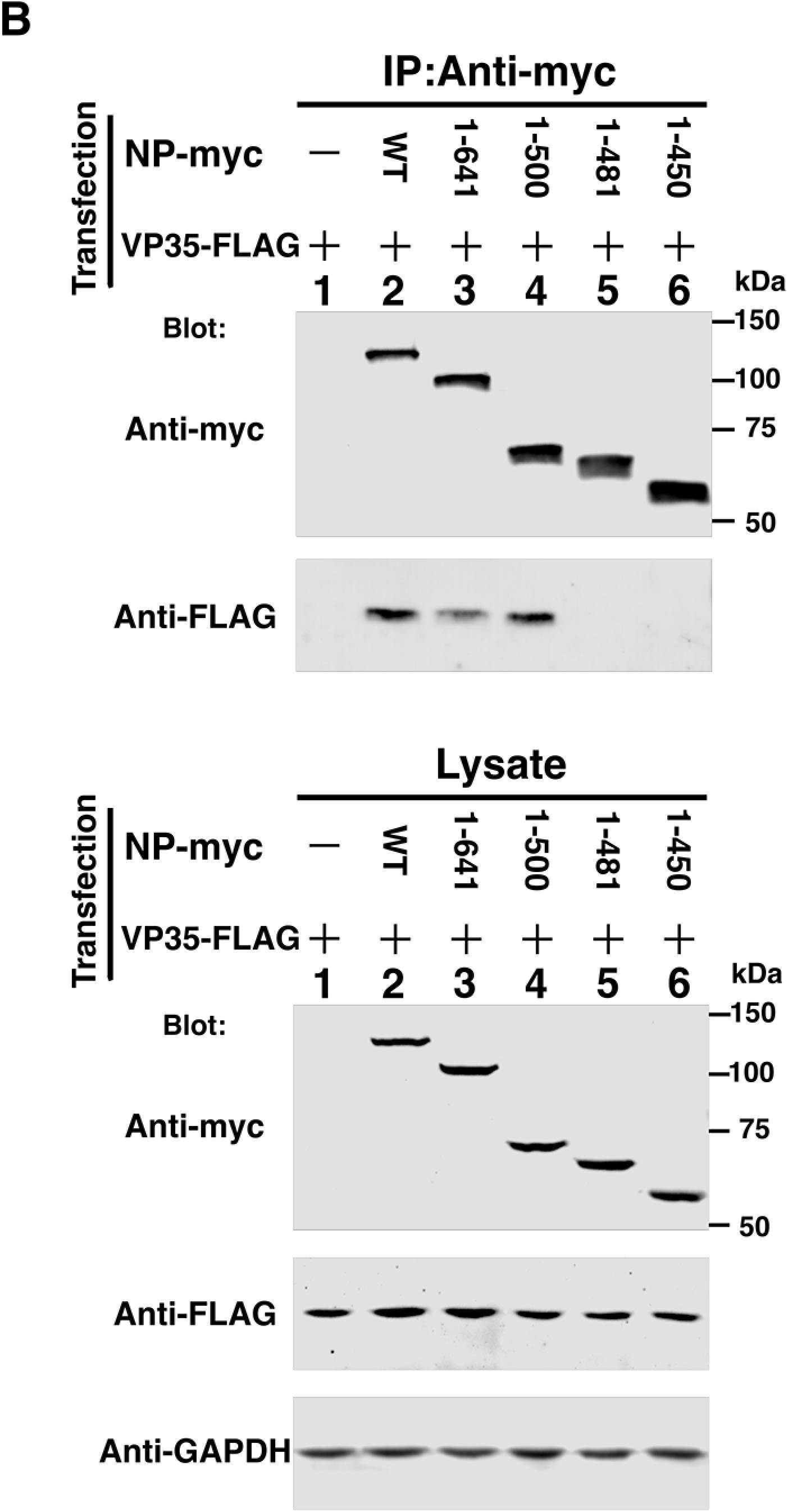
The NP 481-500 region is required for IB formation and association with VP35. A: Localization of co-expressed NP deletion mutants with VP35 in transfected HuH-7 cells. VP35 was detected with anti-VP35 antibody (green) and NP-FLAG proteins were detected with anti-FLAG antibody (red). Nuclei were stained with Hoechst 33342 dye (blue). B: Deletion mutants of myc-tagged NP (NP-myc) and VP35-FLAG were co-expressed in 293T/17 cells and immunoprecipitated with anti-myc antibody. Immunoprecipitated proteins and lysate were subjected to SDS-polyacrylamide gel electrophoresis (PAGE), blotted and detected by anti-myc or anti-FLAG antibodies, as indicated. A GAPDH antibody was used for the loading control.

Next, physical interactions between myc-tagged NP and FLAG-tagged VP35 were examined. As shown in Figure 4B, full length NP, NP(1-641) and NP(1-500) co-immunoprecipitated with VP35, but NP(1-481) and NP(1-450) did not. Identical results were observed using VP35-myc and NP-FLAG constructs (i.e. reversed epitope tags), using immunoprecipitation with anti-myc antibody (data not shown). Thus, our results with co-immunoprecipitation of NP deletion mutants and VP35 were completely aligned with IB co-localization of the identical NP deletion mutants and VP35 (Figure 4A). We conclude that the NP 481-500 region is important for both VP35 association and for IB formation.

### A highly conserved acidic/hydrophobic patch in NP region 481-494 interacts with VP35

Based on our deletion analysis we noticed that the 481-500 region contains a highly conserved acidic/hydrophobic patch, specifically focused within aa 481-494. Sequence conservation is observed across five *ebolavirus* members including EBOV, whose sequence in this region consists solely of acidic and hydrophobic residues (Figure 5A). To examine the importance of this region, three alanine-scanning mutants were constructed to interrogate the possibly redundant activities of the constituent residues. As such, each mutation converted 4 or 5 consecutive residues to alanine stretches (Figure 5A). Initially the mutants were tested in the context of full-length NP expressed alone, but none of them, namely A(482-5), A(485-8) or A(489-93) had any effect on IB formation (Figure 5B). We interpret this result to be due to the presence of an intact NP-Ct in the full-length protein, which provided the redundant IB-formation function. We also tested the localization of NP(1-641) or NP(1-500) with our three alanine scanning mutants. As expected, due to the lack of NP-Ct, these mutants distributed in the cytosol and no IB localization was observed (data not shown). Next, we tested whether the same mutations affect IB formation by NP(1-739), NP(1-641) or NP(1-500) in the presence of VP35. As shown in Figure 5C, all NPs retaining wild-type sequences within the 481-494 region co-localized with VP35 in IBs (top panel). Also, full length NP(1-739) containing alanine scanning mutations localized to IBs, as expected due to the presence of NP-Ct. Importantly however, VP35 overwhelmingly distributed to the cytosol in the presence of NPs containing any of the alanine scanning mutants. There was some minor co-localization of VP35 and NPs with the A(482-5) or A(489-93) mutations, but even in those cases most VP35 clearly spread widely in the cytosol, and not in IBs. This indicates that the NP 481-494 region is required for localization of VP35 in NP-induced IBs. In the case of alanine scanning mutations within NP(1-641) and NP(1-500), both of which lack NP-Ct, no IB co-localization with VP35 was observed. These data clearly suggest physical interaction between the NP 481-494 region and VP35, and that this interaction can establish localization at IBs through VP35-NP complex formation.

**Figure 5.**
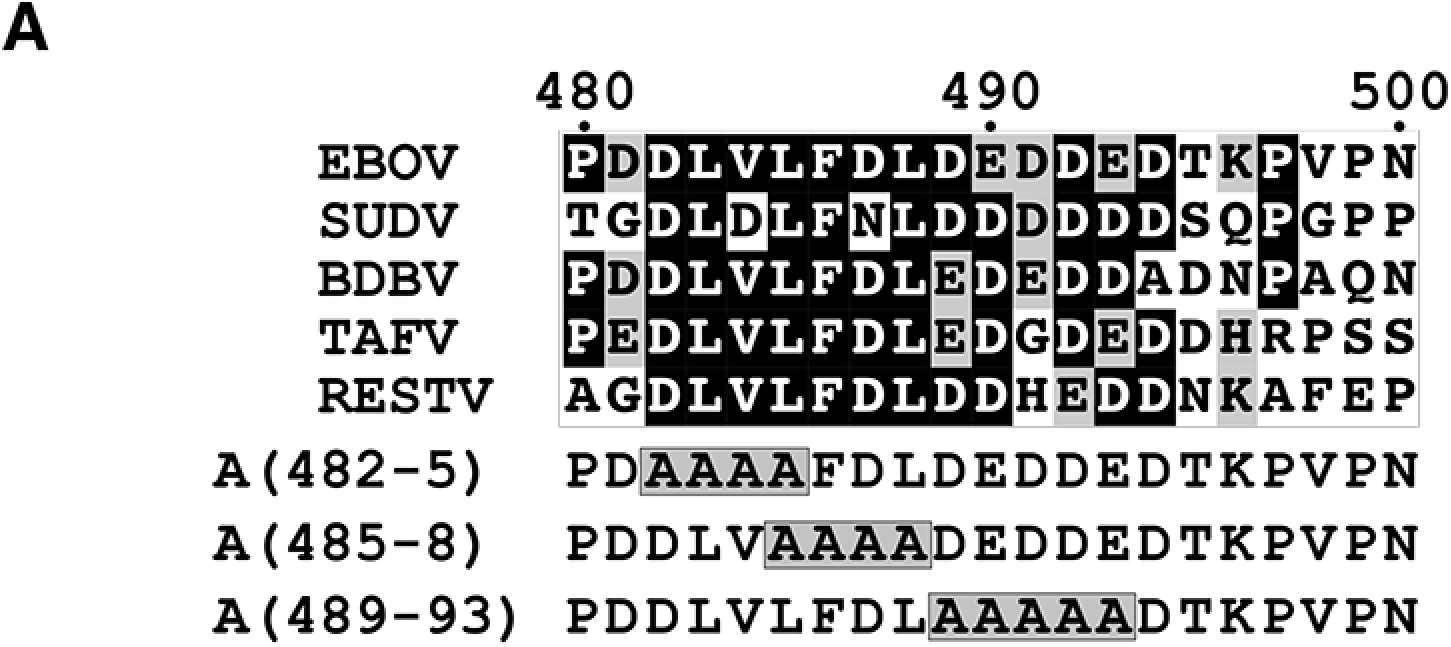
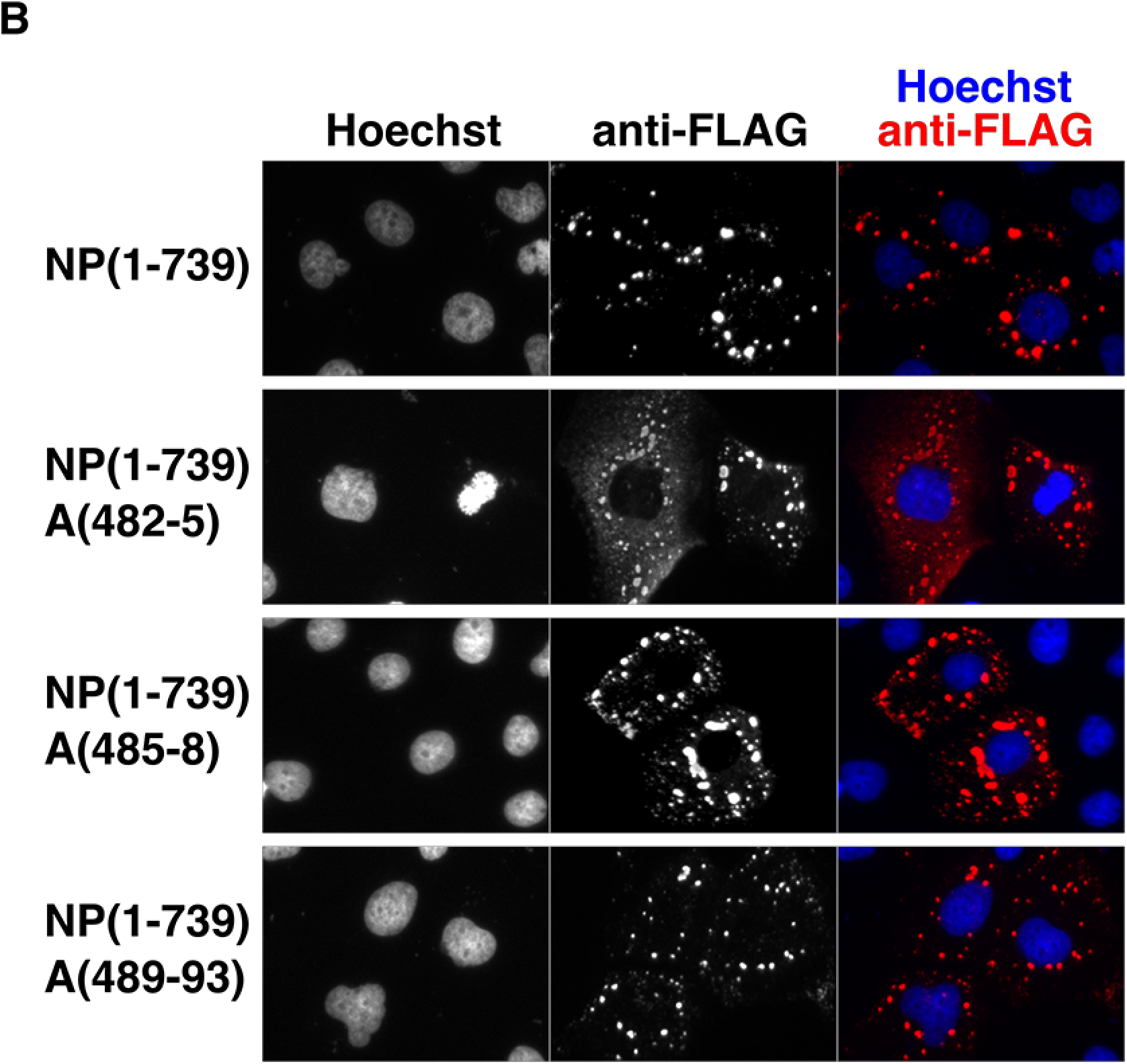
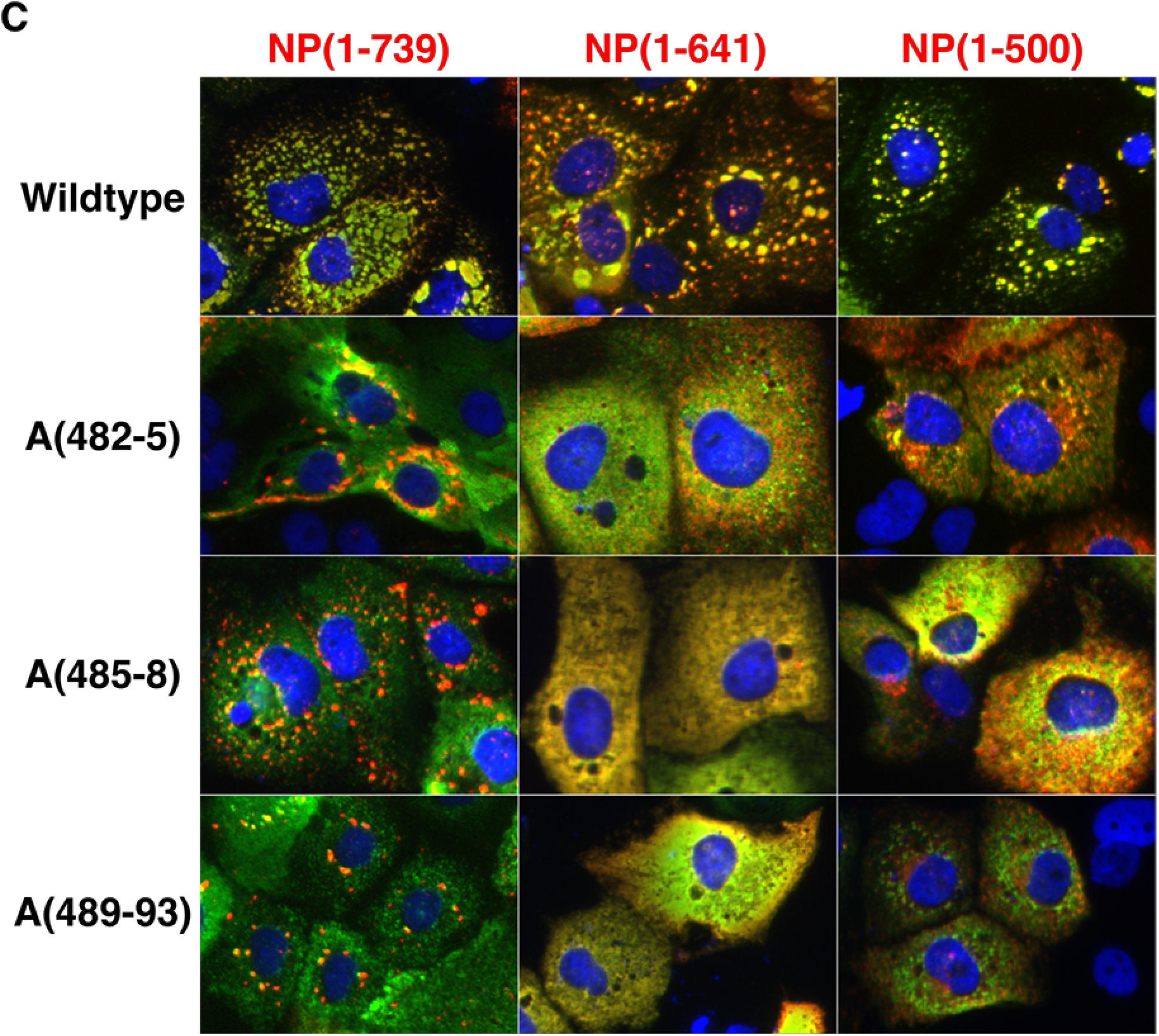
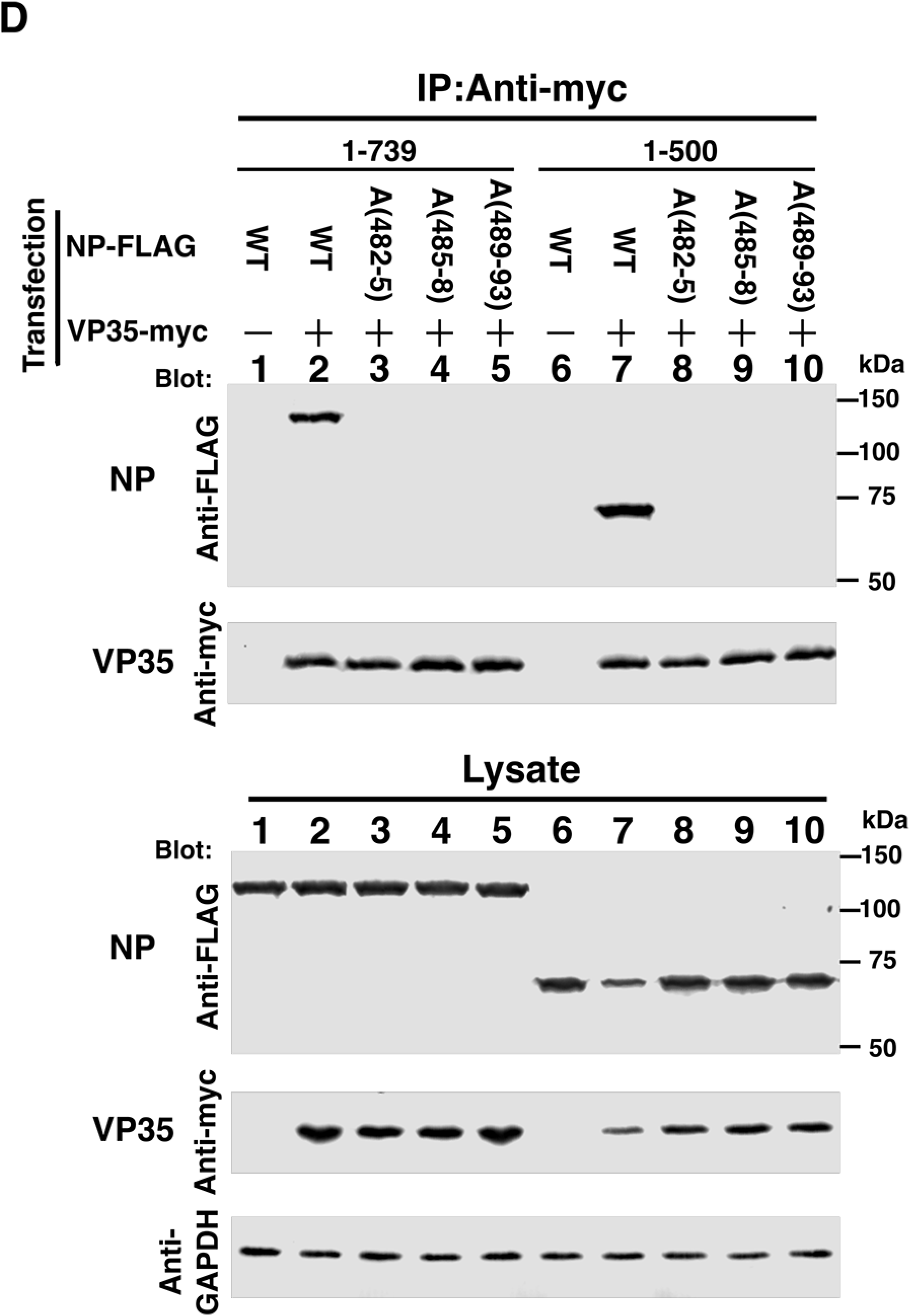
Alanine scanning mutants covering the NP 482-493 region abolish IB formation, co-localization and interaction with VP35. A: Sequence alignment of five *ebolavirus* species members and sequences of alanine scanning mutants within EBOV. Identical residues are shown with white letters on black background, and similar residues are shown with black letters on grey background. Alanine mutations are boxed. B: Immunofluorescence staining of wild-type and alanine scanning mutants of full-length NP. FLAG-tagged NP and mutants were stained with anti-FLAG antibody (red) and Hoechst 33342 dye (blue). C: NP(1-739), NP(1-641) and NP(1-500), or their corresponding alanine scanning mutants were co-expressed with VP35. Cells were stained with anti-FLAG antibody (NP; red) and VP35 was stained with anti-VP35 antibody (green). Nuclei were stained with Hoechst 33342 (blue). Merged green/red fields are shown. D: Immunoprecipitation of co-expressed VP35-myc and wild-type or alanine scanning mutants of FLAG-tagged NP(1-739) or NP(1-500). Cells were co-transfected with VP35-myc and NP-FLAG derivatives, lysed and processed for immunoprecipitation as described in Materials and Methods. A GAPDH antibody was used for the loading control. Top panel: immunoprecipitates were separated by SDS-PAGE, blotted and probed with the indicated antibodies. Bottom panel: Crude lysates were separated by SDS-PAGE, blotted and probed with the indicated antibodies to determine protein expression levels.

To further test this hypothesis, we performed co-immunoprecipitation assays of NP mutants within the 481-494 region (Figure 5D). In the full-length NP(1-739) context, VP35-myc associated with wild-type NP-FLAG, but did not pull down any of the three alanine scanning mutants. The identical result was obtained in the context of NP(1-500) containing each of the alanine scanning mutations. Thus, from co-localization and co-immunoprecipitation experiments, we conclude that the NP 481-500 region, particularly 481-494, is important for the interaction between NP and VP35.

### The IFN inhibitory domain of VP35 is required for NP-VP35 association and co-localization at IBs

Previously, the NPBP (NP binding peptide) of VP35 (aa 20-48) was shown to interact with the NP N-terminal domain (43, 44). NPBP binding is essential for RNA replication and transcription, and it inhibits NP oligomerization and also preserves NP in an RNA-free state (43, 44). We tested whether mutation of the NPBP sequence affects the ability of VP35 to complement deletion of NP-Ct in the formation of IBs. However, mutation of crucial NPBP residues L33D and M34P (43) had no effect on IB formation (not shown). Further deletions of VP35 were constructed to explore the domain(s) responsible for complementing the NP-Ct deletion. As shown in Figure 6A, full length VP35(1-340) co-localized with NP(1-739), NP(1-641), and NP(1-500). Interestingly, VP35(1-219), lacking the interferon inhibitory <u>d</u>omain (IID) located within aa 221-340, failed to co-localize with NP in IBs, or to complement NP-Ct deletion mutants NP(1-641) and NP(1-500). VP35(40-340), which lacks most of the NPBP sequence, did co-localize with all NPs in IBs (full length, NP(1-641) and NP(1-500)). Also VP35(80-340), which completely lacks the NPBP sequence, co-localized with all NPs and localized in IBs.

**Figure 6.**
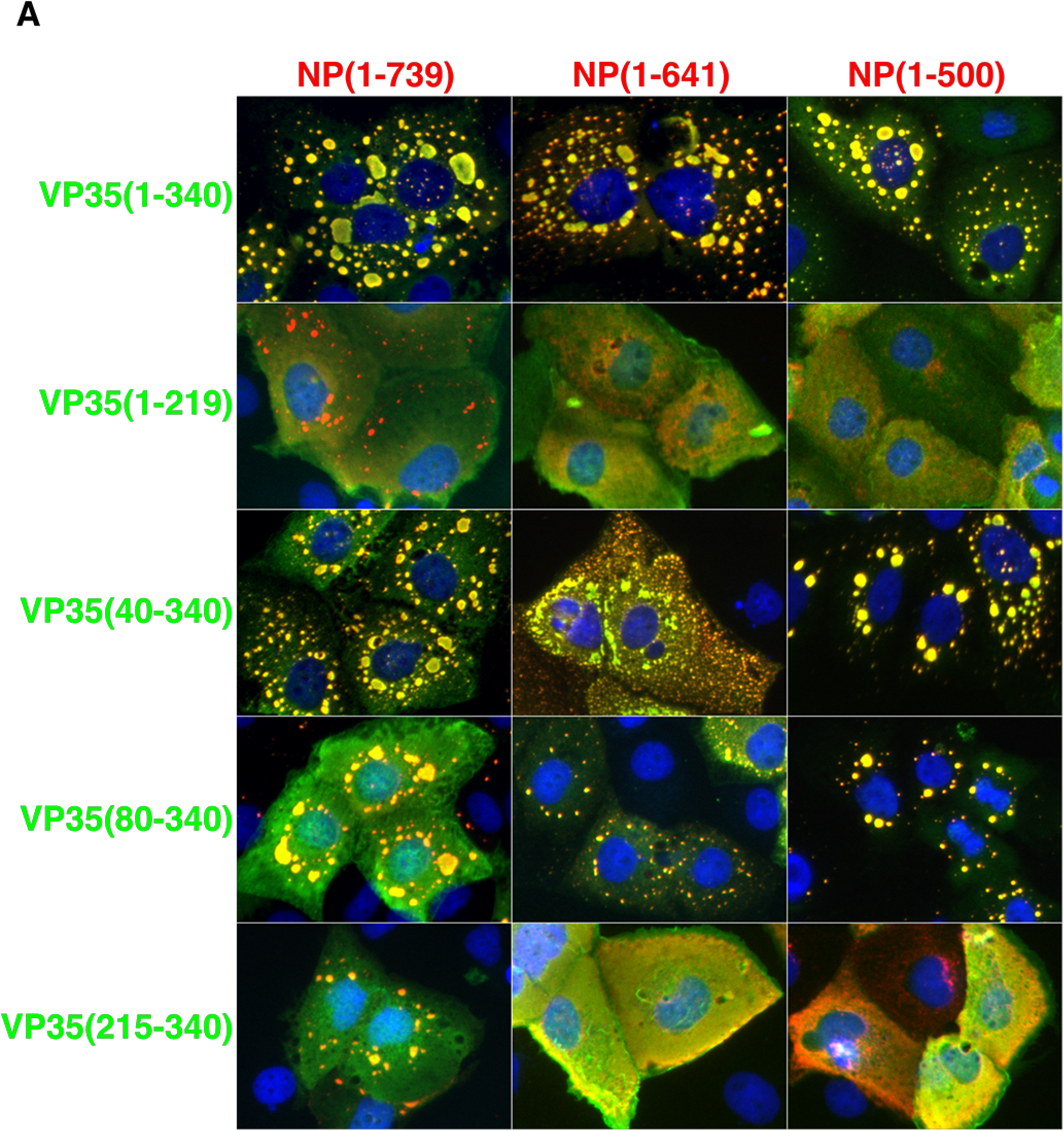
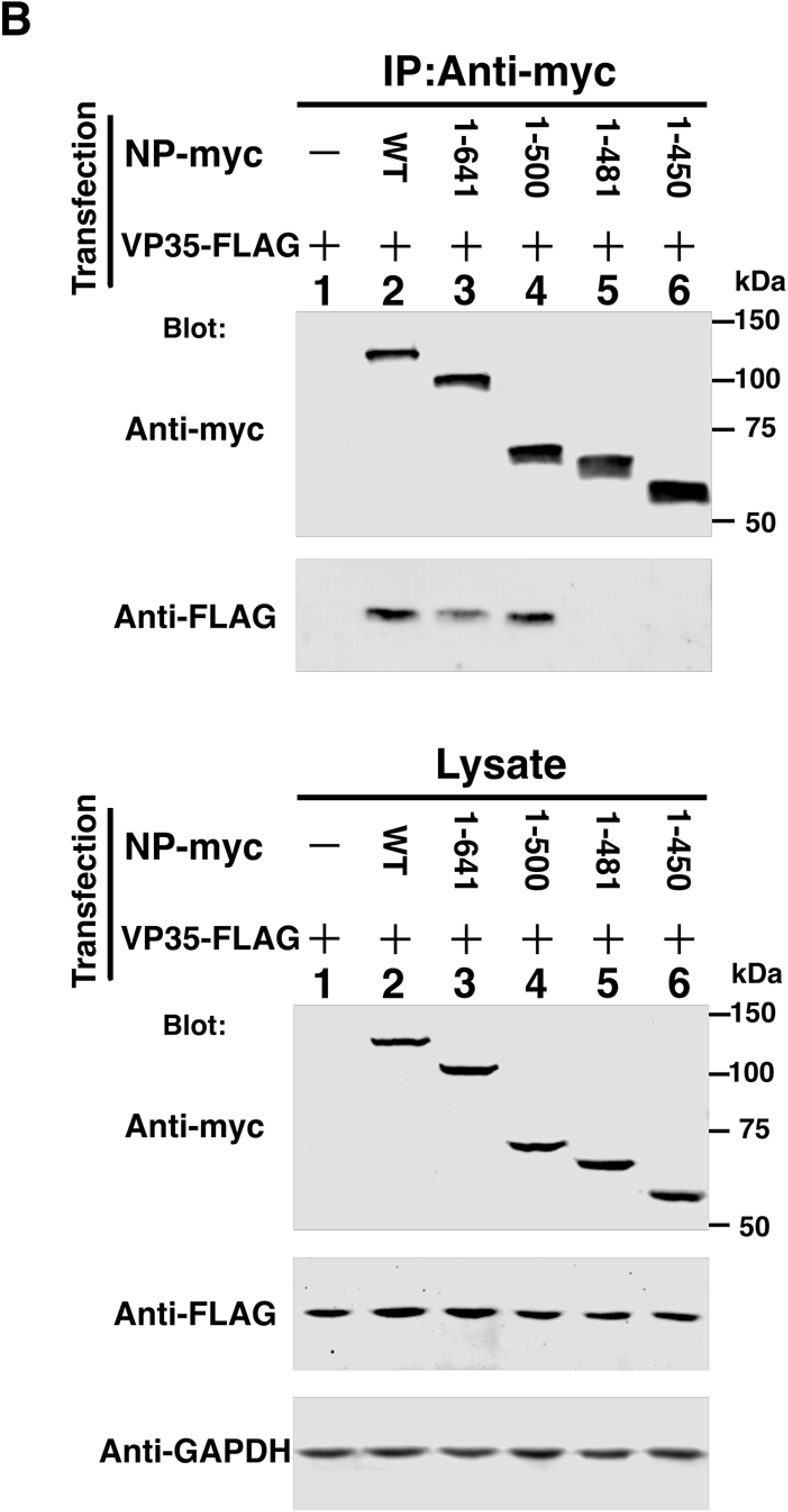
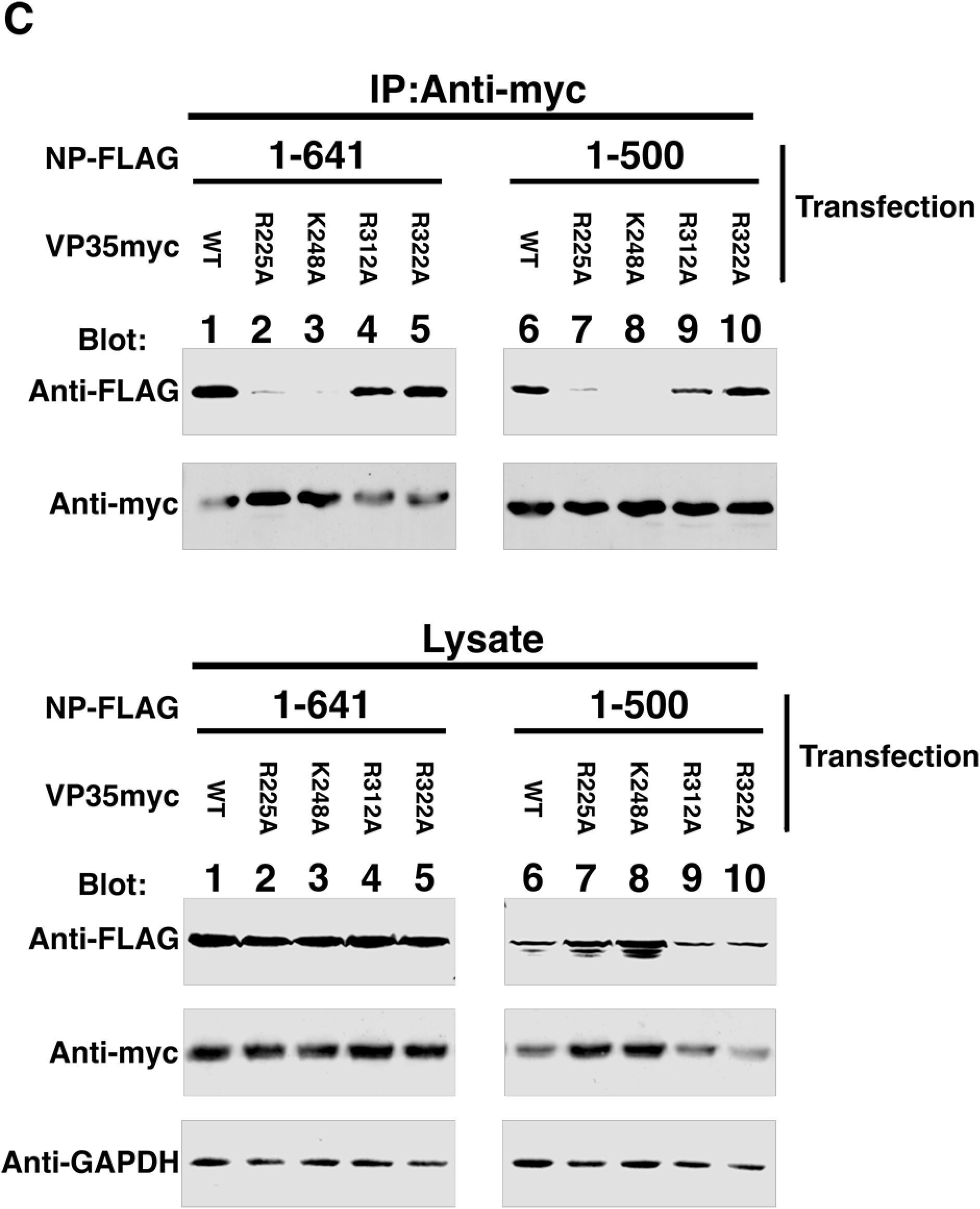
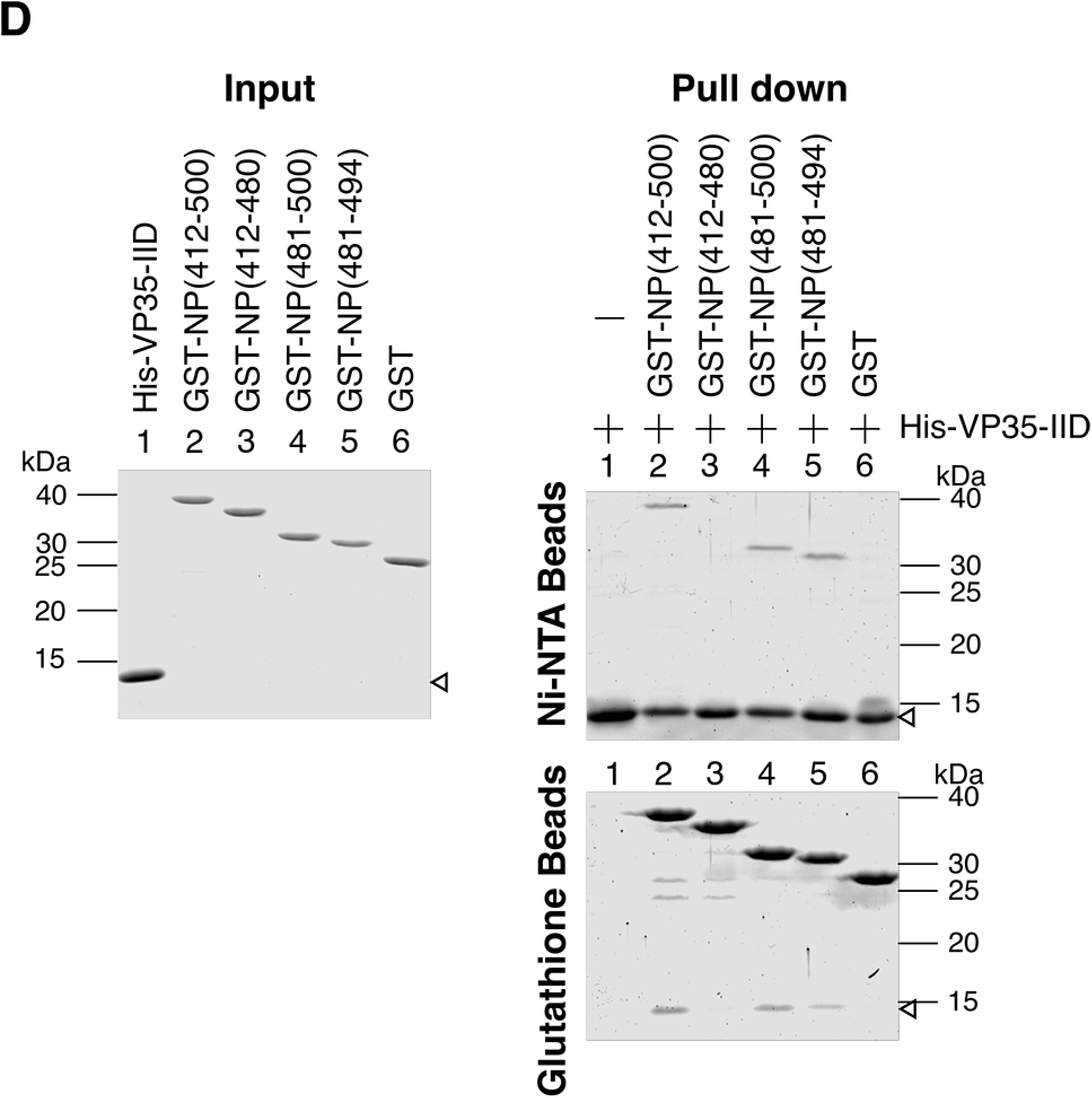
VP35 IID associates with the NP (481-500) region. A: Localization of deletion mutants of VP35 and NP. Wild-type and deletion mutants of NP were co-transfected with VP35 wild-type and deletion mutants in HuH-7 cells. 48 hours after transfection, cells were stained with anti-FLAG (NP, red) and anti-myc (VP35, green). Merged green/red fields are shown. B: NP(1-500) and related mutants as indicated were co-transfected with VP35-myc wild-type and deletion mutants in 293T/17 cells and cells were lysed 48-50 hours after transfection. An anti-myc antibody was used for immunoprecipitation and co-immunoprecipitated proteins were analyzed by western blotting. GAPDH is the loading control of the lysate. Top panel: immunoprecipitates were separated by SDS-PAGE, blotted and probed with the indicated antibodies. The left and right sides of the data shown were sourced from the same image, with several lanes deleted between lanes 5 and 6 of the figure. Bottom panel: Crude lysates were separated by SDS-PAGE, blotted and probed with the indicated antibodies to determine protein expression levels. C: NP(1-500) and NP(1-641) were co-transfected with VP35-myc wild-type or indicated mutants in 293T/17 cells and cells were lysed 48-50 hours after transfection. An anti-myc antibody was used for immunoprecipitation and co-immunoprecipitated proteins were analyzed by western blotting. Top panel: immunoprecipitates were separated by SDS-PAGE, blotted and probed with the indicated antibodies. Bottom panel: Crude lysates were separated by SDS-PAGE, blotted and probed with the indicated antibodies to determine protein expression levels. D: Pull down assay of *E.coli* expressed proteins. GST-NPs with the indicated regions of NP or His-tagged VP35-IID were expressed in *E.coli* and purified using glutathione Sepharose or Ni-NTA agarose. Purified proteins were dialyzed, subjected to SDS-PAGE and stained with CBB (Input panel). Purified proteins were quantified and equal amounts of GST-NPs or GST were mixed with His-VP35-IID protein. After overnight incubation with the indicated beads followed by washing, protein bound to the beads was extracted with 2x sample buffer, and equal volumes of the eluents were subjected to SDS-PAGE and stained with CBB.

As expected, IBs were observed when full length NP(1-739) was co-expressed with VP35(215-340) containing the IID, and the two proteins co-localized. However, the same portion of VP35 didn’t trigger IB formation when co-expressed with either NP(1-641) or NP(1-500) as shown in Figure 6A. This indicates that when NP-Ct is missing, VP35 IID is not sufficient to trigger IB formation, even though both NP(1-641) and NP(1-500) can interact with the IID. Therefore, VP35 sequences between aa 80 and 215 are also required for this function.

To test whether VP35 physically associates with the NP 481-500 region, NP(1-500) and its three alanine scanning mutants were tested for co-immunoprecipitation with VP35 and several of its deletion mutants (Figure 6B). NP(1-500) co-immunoprecipitated with full length VP35 (as also shown in Figure 4B). Consistent with our co-localization data, VP35(1-219), completely lacking the IID, did not associate with NP(1-500). VP35(40-340) and VP35(80-340) did associate with NP(1-500), demonstrating that the NPBP sequence is dispensable for binding of NP(1-500) to VP35. Importantly, VP35(215-340), containing the IID, co-immunoprecipitated with NP(1-500), but not with NP(1-481), NP(1-500/482-5A), NP(1-500/485-8A), or NP(1-500/489-93A), demonstrating that the VP35 IID-NP interaction requires the NP 481-494 region.

Previously it was shown that the “first basic patch” of the VP35 IID (59), binds to NP and is critically important for RNA synthesis in a minigenome assay (45). To test whether these residues are involved in the interaction with NP (481-500), mutants (R225A and K248A), which abolished minigenome activity and binding to NP (45) were tested by pull-down assays (Figure 6C). Two additional VP35 mutants in the “central basic patch” (R312A and R322A) were also tested. These mutants abolish dsRNA binding, reduce suppression of IFN-β promoter activation, but do not affect minigenome activity or NP binding (60). NP(1-641) and NP(1-500), both containing the NP 481-500 region, were examined for binding to all four VP35 mutants. Importantly, mutants R225A and K248A abolished the binding, but mutants R312A and R322A maintained efficient binding. These data are consistent with those of Prins et al. (45) and in addition suggest that residues within the first basic patch of VP35 IID are required for binding to the NP 481-500 region.

To further confirm the interaction between the NP(481-500) region and VP35-IID, GST fusion proteins of NP(412-500) or its shorter derivatives were expressed in *E. coli* and purified, as was His-tagged VP35-IID (Figure 6D “Input”). Next, purified proteins were combined and subjected to pull-down assay. As shown in Figure 6D, we clearly confirmed the association of GST-NP with His-VP35-IID using constructs containing NP(412-500), NP(481-500) and NP(481-494). No binding was observed with either GST alone or with GST-NP(412-480). These data demonstrate that NP(481-494) is sufficient to bind to VP35 IID and identify it as a novel VP35-binding domain. Importantly, for the data presented in Figure 6D, purified proteins were used because this ensured that equivalent amounts of each domain could be strictly compared. To reduce nonspecific binding under these conditions, the protocol was optimized, including elevated salt concentration for higher stringency (see Materials and Methods). The highest salt concentration was 0.5M NaCl, which demonstrated that the binding was resistant to stringent conditions, including detergent. These conditions reduced the amount of recovered protein, but specificity of binding was nonetheless demonstrated.

### The NP 481-500 region is crucial for EBOV RNA synthesis

Given the important role of the NP 481-500 region in IB formation and interaction with VP35, and considering the known functions of NP and VP35 in viral RNA synthesis (31), we asked if the NP 481-500 region is important for reporter activity in p0 cells, which is directly dependent on viral transcription and RNA replication (54-56). Accordingly, trVLP assays of our three alanine-scanning mutants and the corresponding wild-type constructs were performed. As shown in Figure 7, in the context of the full-length NP backbone, all three mutants resulted in strongly reduced reporter activity in p0 cells amounting to 4-12.5% of activity observed in context of wild-type NP. Of note, when p1 cells expressing alanine scanning mutant NPs were infected with wild-type trVLPs from p0 cells, reporter activity in the p1 cells was also strongly reduced, indicating that the mutants were defective in replication of incoming genomes associated with wild-type nucleocapsids (not shown). Keeping in mind that these mutant NPs still localized in IBs due to the presence of NP-Ct (Figure 5B and C), we conclude that the NP-VP35 interaction mediated by the NP 481-500 region is by itself essential for full transcription/RNA replication activity of EBOV. Similar experiments were also performed within the context of the NP(1-641) backbone lacking NP-Ct. Under these conditions, activity of the three alanine scanning mutants was 2.3-2.5% of control NP(1-641) protein in p0 cells, whereas unmutated NP(1-641) fully supported reporter activity in p0 cells (as also shown in Figure 1). Considering that omission of NP altogether from the trVLP assay showed 2.4% the activity of the complete system (“no NP” in Figure 7), clearly these alanine-scanning mutants abolished almost all EBOV RNA synthesis. Taken together, we conclude that NP 481-500 is a crucial region for EBOV replication and importantly, and that its function represents a novel form of regulation at the interface of RNA synthesis and inclusion body dynamics.

**Figure 7.**
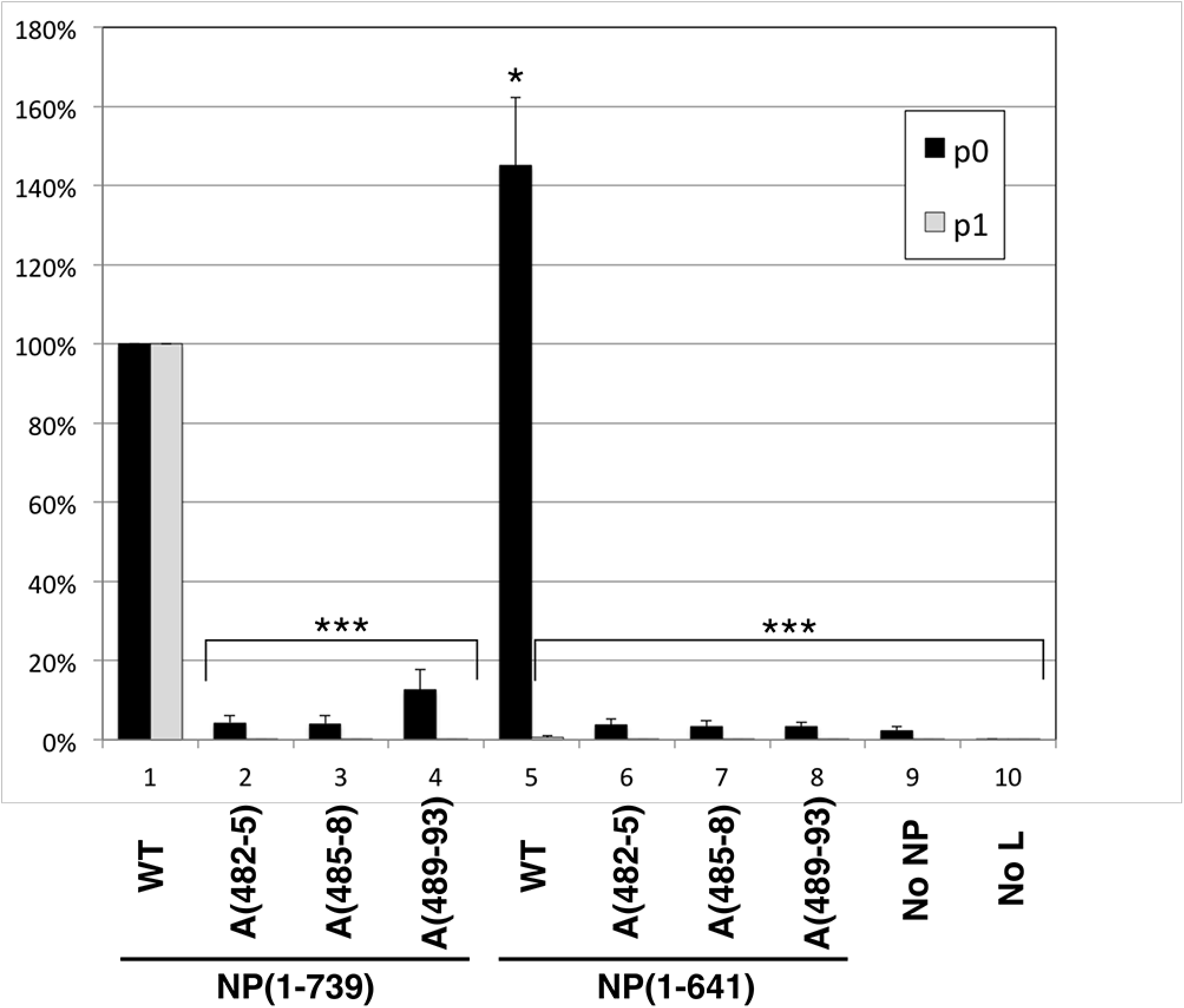
trVLP assay of wildtype and alanine scanning mutants in aa 481-500. For p0 assays, NP(1-739) or NP(1-641) and their alanine scanning mutants, A(482-5), A(485-8) or A(489-93) were co-transfected with other components of the trVLP p0 assay, and luciferase activity of each lysate measured. Wildtype NP (1-739) activity was set at 100%. trVLP assays in the absence of NP (“No NP”) or L (“No L”) were used as negative controls. For p1 assays, recipient cells were transfected with the complete set of trVLP plasmids including wildtype NP, and supernatants from the indicated p0 assay were used for infection. Error bars represent the SD from three independent biological replicates. Asterisk * indicates *p*<0.05, and *** indicates *p*<0.001 to the corresponding NP(1-739) wildtype based on the Student’s t-test.

## Discussion

With this report we have characterized three novel functions of NP, two of which are carried out by NP-Ct, and a third controlled by the newly identified aa 481-500 central domain (CD). Based on our previous structural studies (50-52) we designed a deletion of NP-Ct that abolished production of infectious trVLPs in the trVLP assay, and also abolished IB formation when NP was expressed alone. In the trVLP assay, despite wild-type levels of transcription/replication that were achieved in p0 cells expressing mutant NP(1-641) (Figures 1 and 3A), these cells failed to produce infectious trVLPs. Importantly, since p1 cells in the trVLP assay were pre-loaded with wild-type NP, our results clearly demonstrate that the supernatants from p0 cells contained no infectious trVLPs whose replication would have been supported in p1 cells by the wild-type protein. Further analysis revealed that NP, VP35, VP40 and GP were all detected in similar amounts when comparing trVLPs harvested from NP(1-739) or NP(1-641) expressing cells. Interestingly, even though the protein composition of the two trVLP types were the same, the amount of genomic RNA recovered from the NP(1-641)-associated trVLPs was only 10% of the wild-type (Figure 1E and F). These data clearly suggest that NP-Ct has an important role in genomic RNA incorporation or stability within the VLP nucleocapsids, and as such defines a novel function for this domain. Our finding that NP(1-641) has wild-type transcription/replication activity in p0 cells is consistent with it containing previously identified binding sites for VP35 and VP30 that support transcription/replication, as well as the novel binding site for VP35 described in this work (the CD), in addition to its other well-characterized N-terminal domain functions (aa 1-450) (29, 31, 32).

A second, apparently unrelated function of NP-Ct described here is in the control of IB formation. IB formation accompanies the establishment of viral RNA and nucleocapsid production in the context of viral infection, and also in the context of trVLP or minigenome replication. It is well-established that ectopic expression of NP by itself is sufficient for IB formation (10, 29, 34-37). Our data demonstrate that when NP is expressed alone, IB formation strictly depends on NP-Ct (Figure 2). However, expression of NP-Ct alone is not sufficient to trigger IB formation, suggesting that other regions of the protein are also specifically involved. Since mutant NP(410-739) also fails to make IBs (Figure 2) we conclude that sequences in the N-terminal domain are also required, possibly including the known RNA-binding and/or NP oligomerization functions, to trigger IB formation. Consistent with this, data from Noda et al., using a series of ∼150 aa deletions across NP, showed that only deletion of aa 451-600, which retained an intact NP-Nt and NP-Ct, allowed inclusion body localization (36). The mechanism by which NP-Ct contributes to IB formation is not yet known but may involve interactions with cellular proteins in addition to viral protein interactions or structural regulation of NP. As proposed by Kolesnikova et al, NP helices can be observed by EM in the perinuclear region near ER-bound ribosomes, suggesting that these may be nucleation sites for spatially-directed early viral translation, and that this could explain the emergence of IBs in the perinuclear region (29). NP-Ct could conceivably be involved in this process.

We conclude that the two steps of virus replication supported by NP-Ct (infectious VLP production and IB formation) are mechanistically distinct, because in cells that expressed NP(1-641) plus all other trVLP components intact IBs and transcription/replication were observed (Figure 3), even though these cells failed to produce infectious trVLPs in p0 supernatants (Figure 1). This indicates that failure of NP(1-641) to support infectious trVLP production did not impinge on the process of IB formation. It is interesting, but not necessarily surprising, that the two NP-Ct functions appear mechanistically distinct. The possible roles of NP-Ct in nucleocapsid assembly or overall viral assembly, leading to production of infectious particles, would be expected to involve, at least in part, NP as a stable component of the intact nucleocapsid within newly generating virions. This would likely be distinct from its role in IB formation, which seems to occur in coordination with newly forming or transcriptionally active nucleocapsids, particularly in light of our data presented here, demonstrating that IB formation and RNA synthesis are both controlled by the NP CD. Indeed, in their cryo-electron tomography studies of MARV and EBOV nucleocapsids from intact virus, Bharat et al. (2011) and Bharat et al. (2012) concluded that periodic outward facing protrusions of the nucleocapsid contain the C-terminal region of NP, in addition to VP24 and VP35 (61, 62), which is consistent with a structural role for the C-terminal region in building or stabilizing new virions. In these studies, the NP C-terminal region was more broadly defined than the focused NP-Ct characterized here, but nonetheless NP-Ct might be expected to reside in the outward facing protrusions of the nucleocapsid and therefore be available to be involved in productive virus assembly, including possible contacts with VP40 to achieve proper assembly (36). Additionally, as we demonstrated in Figure 1, NP-Ct is required for viral genomic RNA to be found in isolated VLPs.

Importantly, we found that loss of NP-Ct in deletion NP(1-641) was efficiently complemented by VP35 expression in the formation of IBs, and this correlated with the physical association of NP(1-641) with VP35. Using a combination of deletion and alanine scanning mutations, we identified a novel region of NP (NP 481-500) that interacts with the interferon inhibitory domain (IID) of VP35 and is required for IB formation and RNA synthesis in the trVLP assay. We termed this region the NP central domain based on its location between NP-Nt and NP-Ct. We demonstrated that the VP35 IID is sufficient to bind to NP derivatives containing the CD, including a minimal GST-NP(481-494) fusion protein. The NP 481-494 sequence is highly conserved among five ebolaviruses, and in EBOV it consists of only acidic and hydrophobic residues (Figure 5A). It was previously shown that VP35 IID binds to NP via basic patch residues in VP35, specifically involving R225, K248 and K251 of VP35, and that mutations in those residues abolished the interaction of VP35 IID with NP (45). In this study, we identified the region of NP responsible for VP35-IID binding. It is likely that the VP35 basic residues bind to the acidic residues in NP aa 481-494, because we observed that NP(1-500), which retains the CD, fails to bind VP35 derivatives with mutations in the first basic patch. Also, Prins et al. found that basic patch mutants R225A and K248A abolished an interaction between VP35 and NP (44), which is consistent with our data. Because inhibition of the VP35-CD interaction severely affects trVLP activity, inhibition of this interaction could be a good target for small molecule inhibition. Indeed, compounds that inhibit VP35-NP interactions have been identified (63). Our study also raises the question of why there are apparently redundant functions encoded by EBOV to control IB formation, as revealed by complementation of the NP-Ct deletion by VP35 IID. Our finding clearly suggests that, even though VP35 supports IB formation in the absence of NP-Ct, fully functional IB formation likely requires both VP35 and NP-Ct. Tools are not currently available to distinguish among potentially different “stages” of IB function during infection, but these will be interesting to establish. IBs are turning out to be complex structures composed of viral and cellular components and we speculate that there are interesting, possibly distinct roles for both the NP-Ct and the VP35 binding domain of NP in this process, and that IBs may display different physical and functional properties as infection progresses.

Structural studies by Leung et al. and Kirchdoerfer et al. revealed the binding of a peptide derived from VP35 to the N-terminal domain of NP (43, 44). Binding of the VP35 peptide regulates NP-RNA interactions as well as NP oligomerization, both key requirements for productive RNA synthesis. Moreover the NP region targeted by the peptide, which crucially involves NP residues R240, K248, and D252, is required for maximal RNA synthesis in minigenome assays (44). These findings are the basis for models in which the binding of the VP35 peptide to the N-terminal domain of NP ensures a monomeric and RNA-free state of NP in preparation for productive replication of viral RNA (43, 44). To this picture we have added the dual roles of the NP CD in controlling both RNA synthesis and IB formation. Our data demonstrate that the CD is responsible for interacting directly with sequences within the IID of VP35 and is also crucially required for RNA synthesis and IB formation. Interestingly, this NP-VP35 interaction efficiently complements the defect in IB formation exhibited by the NP-Ct deletion mutant NP(1-641) when it is expressed alone (Figure 3). In our experiments, mutation or deletion of residues within the NP-binding VP35 peptide responsible for controlling RNA binding and NP oligomerization (43, 44) had no effect on the ability of VP35 to complement deletion of NP-Ct or to localize to inclusion bodies (Figure 6 and accompanying text), clearly indicating that the two NP-binding functions of VP35 are separate. This raises interesting questions about the relationship between these two functions of VP35 in supporting RNA synthesis and nucleocapsid assembly. The fact that VP35 interacts directly with NP at two distinct sites could influence the affinity of the individual interactions or the avidity of the overall interaction. One interesting possibility is that the interaction of VP35 IID with the NP CD might prime or enhance the NPBP-NP interaction due to structural influences, thus allowing regulation of NP oligomerization and NP-RNA binding. Figure 8 illustrates the two binding sites within NP for VP35, as defined in this work and by others for the NPBP region of VP35 (43, 44). In addition, Figure 8 illustrates the regions shown here involved in control of IB formation and production of infectious trVLPs. Further work will be required to understand these interesting relationships.

**Figure 8.**
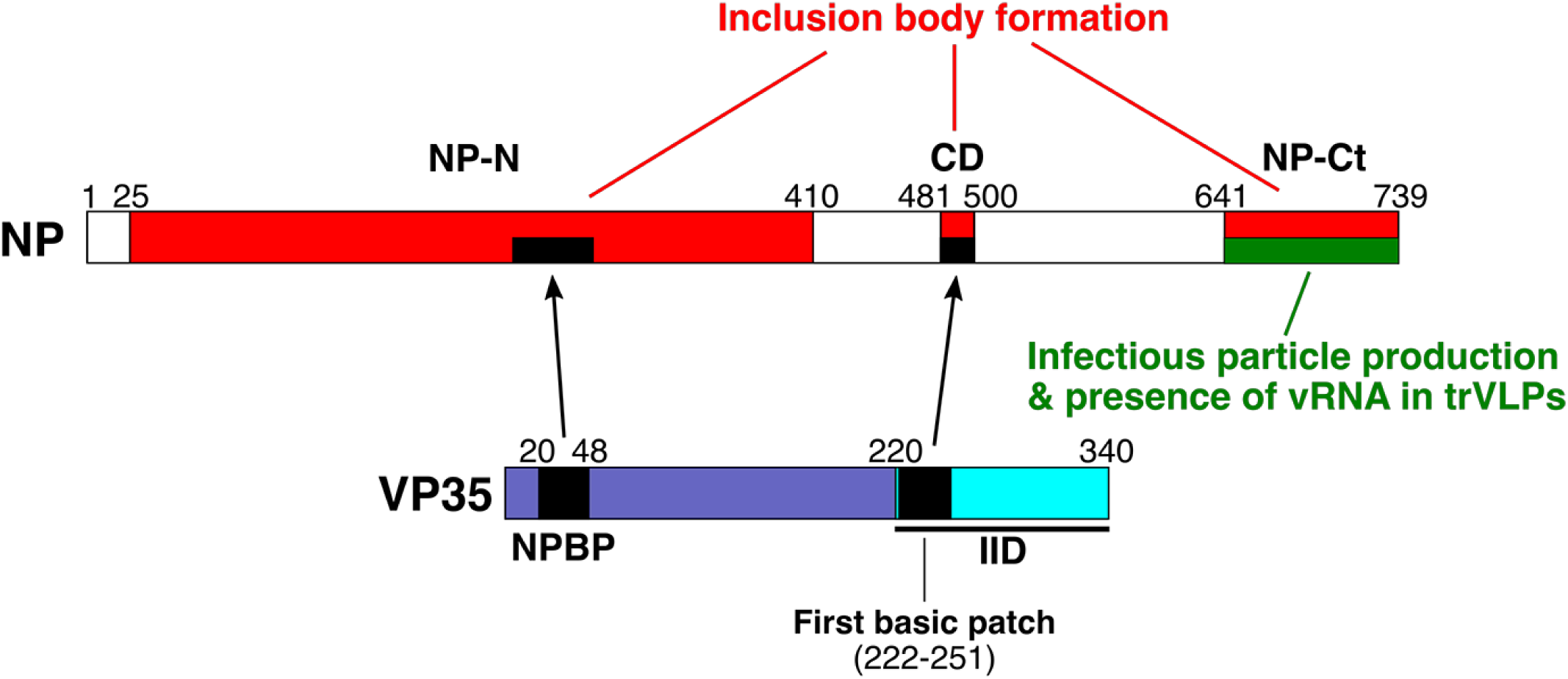
Identified NP functions. Full-length NP and VP35 proteins are illustrated. NP-N and NP-Ct cover aa 1-412 and 641-739 respectively, as defined in (51). The central domain (CD) spans aa 481-500. Red boxes: Regions required for IB formation. The large region spanning aa 25-410 is based on data shown in Figure 2B. The requirement for NP-Ct applies to NP-induced IBs, and when VP35 is co-expressed, NP-Ct is not required, as described in the text. Likewise, when NP-N and NP-Ct are present, mutation of the CD does not abolish IB formation; therefore NP-Ct and CD complement each other. Green box: Region required for production of infectious trVLPs and incorporation/retention of viral RNA in purified trVLPs. For NP and VP35, black regions indicate NPBP and its corresponding binding region within NP, and the VP35 first basic patch as defined in (45, 60), and its corresponding binding region, the CD. See references (43, 44) for definition of NP region bound by NPBP. See Discussion for possible functional relationships between the two regions of VP35 that interact with NP.

Some common themes have emerged from the study of inclusion bodies generated by pathogenic negative strand RNA viruses. These include the housing of RNA synthesis machinery, protection from innate immune mechanisms and association with certain host proteins (although there is not yet strong commonality among the different viruses regarding the identity of these proteins or their function in replication). In some cases there is good evidence that the structure of IBs is achieved by liquid phase separation (26, 27), which is mediated by the activity of specific viral proteins. For EBOV, NP has a central role in IB formation, but there is no evidence so far of a protein shell, phase separation or other process that distinguishes inside from outside. Rather, EM analysis indicates that IBs contain an accumulation of arrays of growing or fully assembled nucleocapsids when cells are infected with live virus or transfected with various nucleocapsid components including NP (10, 42). Also, several groups have reported co-localization with cellular proteins, strongly suggesting that IBs are more than simply accumulations of nucleocapsids that serve as RNA synthesis factories (19, 20, 24). Regardless, whereas the molecular basis for IB formation remains a mystery overall, our results showing the involvement of NP-Ct and the NP CD support a model wherein the earliest steps of IB formation are controlled by NP and its interaction with VP35. However, despite the fact that a small domain of NP is sufficient for direct binding to VP35, and that this interaction controls both IB formation and RNA synthesis, we do not yet know if there is a cause-and-effect relationship between IB formation and RNA synthesis, or whether these two processes are concerted yet mechanistically independent. Clearly, RNA synthesis is not a prerequisite for IB formation. One striking example of this is when all components of the trVLP system are expressed in p0 cells with the exception of the L polymerase, and this results in robust IB formation but no transcription/replication (Figures 1 and 3A). This situation also occurs when full length NPs containing CD mutations are included in the trVLP assay: again, IBs are formed yet transcription/replication is highly defective (Figures 5C and 7). In this case however, the IBs do not contain VP35 due to mutation of the CD (Figure 5C and 6B). These results are consistent with two models. In one model, IB formation and transcription/replication are independent processes, and both are dependent on the interaction between NP CD and VP35. In the second model, transcription/replication strictly depends on IB formation, which depends on VP35 binding to NP. Further investigation of the molecular components of IBs (including the makeup of IBs at different stages of virus replication) will be required to fully distinguish among IB functional models.

## Materials and Methods

### Plasmids

All plasmids for the trVLP assay, i.e. pCAGGS-NP, pCAGGS-VP35, pCAGGS-VP30, pCAGGS-L, p4cis-vRNA-RLuc, pCAGGS-T7, pCAGGS-Tim1, and pCAGGS-luc2, were described previously (64). pCAGGS-NP-FLAG and pCAGGS-VP35myc were made from pCAGGS-NP and pCAGGS-VP35 (55), respectively. All deletions and mutants of pCAGGS-NP-FLAG and pCAGGS-VP35myc were made by using standard PCR and cloning methods. Synthesized sequence of VP35 IID (aa 215-340) was cloned in pET45b (Novagen) to make His-tagged VP35 IID. PCR amplified fragments of the NP(412-500) or shorter fragments from pCAGGS-NP were cloned into pGEX-6P-2 (GE Healthcare). All sequences of the cloned or mutated DNA regions of the plasmids were validated by Sanger DNA sequencing.

### Cells and transfection

293T/17 cells and HuH-7 cells were grown in the media indicated in (65) and (54) respectively. Both cell lines were transfected using TransiT-LT1 (Mirus) according to Biedenkopf et al. (54) except reverse transfection was performed.

### Immunoprecipitation

Forty to forty-eight hours after the transfection, 293T/17 cells were washed once with PBS and lysed in lysis buffer A (50 mM Tris-HCl pH 7.8, 1% NP-40, 150 mM NaCl, 1 mM EDTA) with Pierce Protease Inhibitor Mini Tablets, EDTA-free (Thermo Scientific). The lysates were cleared by centrifugation at 14,000 RPM for 10 min at 4°C. Protein concentration was measured by BCA protein assay kit (Thermo Scientific). Identical amounts of protein in lysates were subjected to immunoprecipitation. Myc-tagged proteins were immunoprecipitated using mouse anti-myc monoclonal antibody (Cell signaling) and rProtein A agarose (Genesee). Identical amounts of lysates were incubated with antibody overnight, then rProtein A agarose was added and incubated for 1 hr. Agarose beads were washed with lysis buffer A 4 times. Proteins bound to the beads were eluted with 2x Laemmli sample buffer. Equal volumes of eluents were subjected to SDS-PAGE.

### Western blotting

Samples were prepared 40-48 hours after transfection by lysing the cells in 60 mM Tris pH 6.8 and 2% SDS. After harvesting the lysates, they were sonicated briefly, and measured for protein concentration. Identical amounts of protein samples were subjected to SDS-PAGE and transferred to Immobilon-FL membrane (Millipore). Immunoprecipitated samples were subjected to SDS-PAGE and transferred to Immobilon-FL membrane (Millipore). FLAG-tagged and myc-tagged proteins were detected with rabbit anti-DYKDDDDK (FLAG) monoclonal antibody and rabbit anti-myc monoclonal antibody (Cell signaling), respectively. For western blotting of trVLPs, the following antibodies were used: rabbit anti-NP antibody (Genetex), mouse monoclonal anti-VP35 (Kerafast), mouse monoclonal anti-GP, H3C8 (a gift from Dr. Judy White), and rabbit anti-VP40 (IBT Bioservices). GAPDH mouse monoclonal antibody (Millipore) was used for the loading control. Goat anti-rabbit antibody conjugated with IRDye680RD and goat anti-mouse antibody conjugated with IRDye800CW were used as secondary antibodies. Odyssey CLx (Li-Cor) was used to scan the membranes.

### Immunofluorescence staining

Expression plasmids of the proteins indicated in the figures were transfected into HuH-7 cells and plated on coverslips in 6-well plates. “All trVLP” is the combination of following; pCAGGS expression plasmids encoding NP-FLAG or NP(1-641)-FLAG (125 ng/well), VP35 (125 ng/well), VP30 (75 ng/well), L (1000 ng/well), T7-polymerase (250 ng/well), and a p4cis-vRNA-RLuc plasmid (250 ng/well). Forty to Forty-eight hours after the transfection, cells were washed once with PBS and fixed with 4% formaldehyde in PBS. After permeabilization with 0.1% trition X-100 (Sigma-Aldrich) in PBS, cells were blocked with 1x diluted Odyssey Blocking Buffer (PBS) (Li-Cor). Cells were stained with primary antibodies: anti-DYKDDDDK, anti-myc antibody (Cell signaling) and/or mouse anti-VP35 antibody (Kerafast), then stained with secondary antibodies: Alexa Fluor 488 goat anti-mouse IgG (H+L) and Alexa Fluor 594 goat anti-rabbit IgG (H+L). The nuclei were stained with Hoechst 33342 dye. Prolong Gold antifade reagent (Invitrogen) was used for mounting.

### trVLP assay

trVLP assays were performed according to Hoenen et al. (55) except that reverse transfection was performed. pCAGGS-NP was replaced with pCAGGS-NP-FLAG or its derivatives in either p0 or p1 cells as indicated in the figures.

### Protein and RNA analyses of isolated trVLPs

Procedures are described in Watt et al (56). with minor modifications. For western blotting of purified trVPLs, 24 ml of cell supernatant (72 hrs after transfection) was harvested and centrifuged twice at 2500 RPM for 10 min to remove cell debris, then concentrated by ultracentrifugation through a 20% sucrose cushion in an SW-32 rotor at 25,000 RPM for 2.5 hrs at 4°C. Pellets were resuspended in 140 μl of PBS and 47 μl of 4x sample buffer was added. Equivalent volumes of samples were subjected to western blot analysis. For RNA quantification, RNA was purified from 280 μl VLP-containing supernatant using a QIAamp viral RNA (vRNA) Mini Kit (Qiagen) according to the manufacturer’s instructions. Sixteen μl of RNA was subjected to a 30-min DNase digest using 2 μl DNase I (Thermo Scientific) in a total volume of 20 μl according to the manufacturer’s instructions. Digested RNA (7.5 μl) was reverse transcribed using Super-Script III reverse transcriptase (Life Technologies) according to the manufacturer’s instructions with the primer 5-CGGACACACAAAAAGAAAGAAG-3. Five μl of the resulting cDNA was amplified by touchdown PCR (10 cycles of annealing at 59 to 54°C for 30 s, followed by 10 cycles of annealing at 54°C for 30 s) using Taq polymerase (New England Biolabs) according to the manufacturer’s instructions and the primers 5-CTTGACATCTCTGAGGCAAC-3 and 5-ATGCAGGGGCAAAGTCATTAG-3. One μl of PCR product was then subjected to standard PCR with Taq polymerase using the primers 5-CGAACCACATGATTGGACCAAG-3 and 5-CTTATCAGACCTCCGCATTAATC-3. Quantification standards were included in each PCR at the stage of the first amplification. Known amounts of minigenome DNA were amplified along with RT-PCR under the same conditions and quantified to establish the linear range of amplification. A linear range of amplification was verified from the standards for all trVLP samples. Intensity of bands after agarose gel electrophoresis was measured using the Molecular Imager XRS system and quantified with Image Lab (Bio-Rad).

### Pull down assay of *E.coli* expressed proteins

His-tagged VP35 IID (VP35 aa 215-340) and GST-NP (aa 412-500) or shorter variants were expressed in the BL21 CodonPlus strain (Agilent Technologies). Cells were lysed in Buffer (0.15) (50 mM Tris-HCl pH 8.0, 0.2 mM EDTA, 150 mM NaCl, 5 mM imidazole, 20% glycerol) with Pierce Protease Inhibitor Mini Tablets, EDTA-free (Thermo Scientific) and 0.1 mg/ml of lysozyme (Sigma-Aldrich) and then the suspension was sonicated. After centrifugation, supernatants were subjected to purification using either HIS-Select HF Nickel Affinity Gel (Sigma-Aldrich) or Glutatione Sepharose 4B (GE healthcare). Purification was confirmed by gel electrophoresis and Coomassie Brilliant Blue (CBB) staining. After dialysis with Slide-A-Lyzer MINI Dialysis Devices, 3.5K MWCO (Thermo Scientific), purified proteins were quantified by BCA protein assay kit (Thermo Scientific). Protein concentration was set to 1μM of His tagged proteins and to 2 μM of GST-tagged proteins for HIS-Select HF Nickel Affinity Gel pull down, and was set to 1 μM of GST tagged proteins and to 2 μM of His-fusion proteins for Glutatione Sepharose 4B pull down. Purified proteins were mixed as indicated in the figure and rotated with either HIS-Select HF Nickel Affinity Gel (Sigma-Aldrich) or Glutatione Sepharose 4B (GE healthcare) overnight. Beads were washed twice with Buffer (0.15) with 0.1% NP-40, once with Buffer (0.35) and Buffer (0.5), those are identical to Buffer (0.15) with 0.1% NP-40 except including 0.35 M and 0.5 M NaCl, respectively. Samples washed once again with Buffer (0.15) containing 0.1% NP-40. Proteins on the beads were eluted with 2 x sample buffer. Identical volumes of the samples were subjected to SDS-PAGE, and stained with CBB.

## Acknowledgements

The work was supported by NIH grant 1R21AI130420 to DAE. We thank Elizabeth Nelson and Prof. Judy White for reagents and helpful discussions.

